# Rapid changes in cholinergic signaling, myelination and thyroid signaling pathway gene expression in amygdala subnuclei in response to social status maintenance and reorganization

**DOI:** 10.1101/2025.03.28.646014

**Authors:** Tyler M Milewski, Won Lee, Köll R Rada, James P Curley

## Abstract

Male CD-1 mice form linear social hierarchies and can rapidly reform them following social reorganization. Through Tag-based sequencing in the medial amygdala (MeA), we identified several genes regulating cholinergic signaling, myelination, and thyroid signaling that rapidly shift expression 70 minutes after animals change social status. Here, we further characterize the expression patterns of individual genes within these pathways in both stable and reorganized hierarchies. We find that genes related to cholinergic signaling show higher expression in the MeA of dominant males in stable hierarchies as well as when reestablishing dominance in reorganized hierarchies. Dominant males also show higher levels of myelination related genes than socially descending males when reestablishing their social status during social reorganization but less so in stable groups. Conversely, thyroid signaling genes show higher expression in the MeA in subordinate males and previously dominant males who are socially descending. Using RNAscope, we were able to demonstrate broadly similar patterns of gene expression immediately following social reorganization across the MeA, basolateral, and central amygdala for 7 genes of interest (*chat*, *slc5a7*, *ache*, *mbp*, *mog*, *crym*, *mybpc1*). High levels of co-expression of cholinergic signaling and myelination gene expression in dominant males suggest that these processes work together to promote resilience to the social challenge and promote dominance. In summary, we demonstrate that rapid changes in amygdala gene expression in each pathway are associated with the formation and maintenance of dominance and subordinate social status in stable and reorganized environments.

## Introduction

Across the animal kingdom, vertebrates have evolved to be social and live in complex groups. For species living in social environments, the ability to competently interact with others is essential for an animal’s survival and reproductive fitness (Milewski et al., 2022; Taborsky & Oliveira, 2012). To respond with appropriate social behaviors to particular social contexts, animals must assess their external environment and current physiological state and integrate this information with contextual and social memories (Anderson, 2016; Chen & Hong, 2018; O’Connell & Hofmann, 2011). Social dominance hierarchies are a common form of social organization that require animals to continuously make social decisions related to their relative social rank compared to other individuals (Chase, 1982; Chen & Hong, 2018). For example, when establishing and maintaining social hierarchies, individuals must plastically modulate their behavior based on past experiences with each conspecific in their group to dynamically adjust the levels of dominance or subordinate behaviors towards animals of relatively lower or higher rank. Social hierarchies tend to be stable in controlled conditions with relatively few changes in rank among group members. However, when the social context of the hierarchy changes, such as when relatively more dominant individuals die or leave the group, or there is a change in group composition through immigration or fusion of groups, individuals must flexibly change their behavior to adjust to the new social environment appropriately (Curley, 2016; Desjardins et al., 2012; Maruska & Fernald, 2010, 2013; Snyder-Mackler et al., 2016; So et al., 2015; Williamson et al., 2017). Experimentally, the rapid reestablishment of social hierarchies and ranks has been shown through the reorganization of individuals from stable social hierarchies into novel social groups (e.g., African cichlid fish Friesen et al., 2022; Maruska, 2015; guppies - Chase et al., 2002 PNAS, monk parakeets - Hobson & DeDeo, 2015, rhesus macaques - Snyder-Mackler et al., 2016). Further, subordinate male mice and African cichlid fish both inhibit their aggression when a dominant alpha individual is present (Curley, 2016; Desjardins et al., 2012) but can rapidly ascend in social rank upon the experimental removal of an alpha individual from the group (Friesen et al., 2022; Maruska, 2014; Maruska & Fernald, 2013; Williamson et al., 2017; Williamson et al., 2019). Changes in these social behaviors are governed by brain circuits involved in social decision-making that are conserved across social vertebrates and constitute the Social Decision Making Network (SDMN) (O’Connell & Hofmann, 2011). For example, increases in immediate early gene (IEG) activity throughout the SDMN occur as mice and cichlids socially ascend or descend in social status (Maruska, 2015; Williamson et al., 2019), suggesting that coordinated changes in brain plasticity are associated with changing social status.

We have recently demonstrated that outbred CD-1 male mice form new social hierarchies within minutes when rearranged into novel social groups with three other unfamiliar mice of equivalent social status (Milewski et al., 2025). In such groups of four males, individual mice either socially ascend, descend, or maintain their original social rank. We identified large changes in the transcriptome of rising and descending males in the medial amygdala (MeA) when changing social ranks with the greatest shifts in gene expression being found in males were originally dominant when placed into the new social hierarchy (Milewski et al., 2025). The MeA is a central hub of the SDMN and plays a critical role in a wide range of social behaviors, integrating sensory and behavioral pathways (O’Connell & Hofmann, 2011; Petrulis, 2020; Raam & Hong, 2021). The MeA consists of four anatomical subdivisions: anterodorsal (MeAad), anteroventral (MeAav), posterodorsal (MeApd), and posteroventral (MeApv). The MeA receives input directly from olfactory processing areas and projects outputs to several downstream nuclei that regulate behavioral output, including dominant-subordinate relationships in a social hierarchy (Hong et al., 2014; Raam & Hong, 2021). Specifically, in the MeA 70 minutes after dominant males socially descended, we observe large and significant reductions in the expression of genes related to cholinergic signaling including choline o-acetyltransferase (*chat*), cholinergic receptor muscarinic 2 (*chrm2*), and three solute carrier genes (*slc18a3*, *slc10a4*, *slc5a7*) representing the formation and processing of acetylcholine (Milewski et al., 2025).

Socially descending males also show significant decreases in the expression of genes related to the structure and development of myelin such as myelin and lymphocyte protein (*mal*), myelin-associated glycoprotein (*mag*), myelin basic protein (*mbp*), myelin oligodendrocyte glycoprotein (*mog*), and myelin-associated oligodendrocyte basic protein (*mobp*) when compared to dominants who maintained their social status. Additionally, socially descending males have heightened expression of genes related to thyroid signaling (crystallin mu: *crym* and thyrotropin-releasing hormone receptor: *trhr*) relative to dominant males who maintain their dominant status (Milewski et al., 2025).

These three pathways (cholinergic signaling, myelination, thyroid signaling) have been previously implicated in regulating behavioral responses to social challenges. Cholinergic signaling in the forebrain promotes social memory formation (Okada et al., 2021) and is modulated by social defeat stress (Mineur et al., 2013, 2016, 2018). In the basolateral amygdala, increased levels of acetylcholine and nicotine receptors have been associated with depressive-like phenotype in mice after social defeat stress, with local infusions of a nicotinic receptor antagonist decreasing depressive-like behaviors (Mineur et al., 2016). Muscarinic and nicotinic receptors promote and inhibit aggression, respectively (Bell et al., 2009). Elevated levels of acetylcholine (ACh) and activation of muscarinic M1 receptors (M1) in the prelimbic cortex (PL) are associated with increased dominance behavior in social hierarchies (Chen et al., 2023). The remodeling of myelination through activity-dependent processes is a key component of neural plasticity and learning following changes in social environments (Chang et al., 2016; Fields, 2015; Purger et al., 2016). Chronic social defeat stress leads to long-term reductions in myelination across many areas of the SDMN (Bonnefil et al., 2019; Lehmann et al., 2017), but can also lead to increases in myelination in a brain region-specific manner (Poggi et al., 2022). Social stressors are associated with reduced expression of oligodendrocyte genes encoding myelin proteins (Cathomas et al., 2019) in the central and basolateral amygdala, whereas increases in oligodendrocyte progenitor markers are observed in the amygdala of the mice after being housed in socially enriched environments (Okuda et al., 2009). Thyroid signaling is also associated with neuroplasticity during learning (Raymaekers & Darras, 2017). and regulates fear learning (Bárez-López et al., 2017; Montero-Pedrazuela et al., 2011) and social vigilance (Kwon et al., 2021) in the amygdala specifically. Further, the removal of the thyroid leads to reduced myelin and myelin-related proteins in the brain of developing rats (Balázs et al., 1969), which leads to persistent reduction of choline acetyltransferase in the basal forebrain throughout adulthood (Patel et al., 1987), suggesting that these three systems may be functionally interrelated.

In this study, we used *in situ* hybridization (RNAscope) to spatially assess changes in the expression of genes associated with cholinergic signaling, myelination and thyroid hormone signaling in the MeA and other subnuclei of the amygdala as dominant males socially descend. Using Tag-based RNA Sequencing, we also determined if these changes persisted up to 24 hours later beyond the initial descent or whether they were associated with immediate responses to social descent. Finally, we examined whether these expression changes are specific to transitions in social rank or also exist between dominant and subordinate males living in stable hierarchies. We discover that cholinergic signaling genes are relatively upregulated in dominant individuals regardless of whether they were socially reorganized or not. Genes associated with myelination are expressed most strongly in dominant individuals who maintain their status during a social reorganization and thyroid signaling-related genes are more highly expressed in socially descending and subordinate animals.

## Methods

In **experiment 1**, we examined the expression of genes associated with cholinergic signaling, myelination and thyroid hormone signaling in the MeA of male mice either undergoing social hierarchy reorganization or living in stable social hierarchies. Using Tag-based RNA Sequencing we compared expression of genes in males who socially descended or remained socially dominant in groups of four mice at 70 minutes or 25 hours following hierarchy reorganization. For mice living in socially stable hierarchies, we compared the expression of these genes in the MeA of dominant, sub-dominant and subordinate males from mice living in groups of ten males. In **experiment 2**, we used RNAscope to visualize and quantify the expression of select genes across several amygdaloid nuclei 70 minutes after social hierarchy reorganization in males that maintain social dominance or socially descend in groups of four.

For experiment 1, MeA samples at 25 hours post social hierarchy reorganization were newly sequenced from animals whose behavior was previously published (Milewski et al. 2025). Tag-based RNA sequencing data from MeA samples at 70 minutes post social hierarchy reorganization was reanalyzed from Milewski et al. 2025. MeA samples from males living in stable social hierarchies were newly sequenced from animals whose behavior was previously published (Lee et al., 2022). For experiment 2, we ran new groups of males through our social hierarchy reorganization paradigm to confirm the behavioral findings from our previous study (Milewski et al., 2025). The brains of these males were used for RNAscope analysis. All procedures were conducted with the approval of the Institutional Animal Care and Use Committee of the University of Texas at Austin.

### Experiment 1 – Animals, Housing, and Behavioral Observations

MeA samples for Tag-Sequencing from male mice who were socially reorganized were from animals whose behavior was previously published (Milewski et al., 2025). In brief, male CD-1 mice (N = 208) aged 7-8 weeks were obtained from Charles River Laboratory (Houston, TX, USA) and housed at The University of Texas at Austin. The animal facility was kept on a 12/12 light/dark cycle, with white light at 2300 h and red lights (dark cycle) at 1100 h with constant temperature (21–24 °C) and humidity (30–50%). Upon arrival, mice were marked with a nontoxic animal marker (Stoelting Co., Wood Dale, IL, USA) to display unique identification and randomly placed into 52 groups of four animals for 11 days in two standard rat cages (35.5 x 20.5 x 14cm) covered in pine shaving bedding and connected by Plexiglass tubes (diameter 38 mm) with multiple enrichment objects (**Supplemental Figure 1A**). Standard chow and water were provided *ad libitum*. From group housing day 2 (GD2), live behavioral observations began and were conducted daily during the dark cycle until GD11 by a trained observer, as described previously (So et al., 2015; Williamson et al., 2016). Briefly, all occurrences of agonistic (fighting, chasing, mounting) and subordinate behaviors (freezing, fleeing, subordinate posture) between two individuals were recorded using either an Android device or Apple iPad and directly uploaded to a timestamped Google Drive via a survey link. On average, 45 minutes of live observation per group was conducted daily, averaging 65 agonistic/subordinate behavioral interactions. At the start of GD11, between ZT23.5 and ZT00, all alpha (rank 1) individuals from each group of stable social hierarchies were reorganized (N= 13 cages, 52 males) into a new group of four unfamiliar animals of equal (i.e., alpha) social status. Animals remained in these reorganized groups for 70 minutes (N=7 cages of 4 alpha males each) or 25 hours (N=6 cages of 4 alpha males each), during which live behavioral observations were conducted immediately following social reorganization. Consequently, some alphas remained socially dominant in their new groups, whereas others descended in social ranks either to rank 2, 3, or 4.

MeA samples for Tag-Sequencing from alpha, subdominant, and subordinate male CD-1 mice who were living in stable social hierarchies were from animals whose behavior was previously published (Lee et al., 2022). Male mice (N=110) eight weeks of age were housed in a custom-built vivarium for 14 days, as previously described (Williamson et al., 2016). Briefly, each vivarium consists of an upper level with multiple shelves (total available surface: 36,000 cm2 = 3 floors × 150 cm × 80 cm) and a lower level with five connected nest boxes (2,295 cm2 = 5 cages × 27 cm × 17cm). The total surface of a vivarium is 62,295 cm2 (6,230 cm2 per mouse) and was covered in pine shaving bedding (**Supplemental Figure 1B**). Food and water were provided at the top of the vivarium *ad libitum*, and mice could access all levels of the vivarium through interconnected tubes, nest boxes, and shelves. 11 vivaria were used in total each housing 10 males. Experimenters observed the social behaviors of each group for a minimum of 2 hours a day to obtain hierarchical behavior data. An extensive characterization of the behavior of animals used in this dataset can be found in Lee et al., 2022.

### Experiment 1 – Tag-based RNA Sequencing

Following behavioral observations, brains were flash-frozen in hexane and stored at - 80°C until dissection. Whole brains were sectioned on a cryostat (Lecia Biosystems, Deer Park, IL) at 300µm. Then the medial amygdala (MeA both anteroventral and anterodorsal) was dissected separately using a Stoelting 0.51 mm tissue punch (Stoelting, Wood Dale, IL) and homogenized in 100 μl lysis buffer (Thermo Fisher Scientific, Waltham, MA; MagMax Total RNA isolation kit) with 0.7% beta-mercaptoethanol by vortexing at 3000 rpm speed for 15–20 s. Lysates were incubated at room temperature for 5 min and stored at −80 °C until RNA extraction. After all dissections were completed, lysates were further processed on KingFisher Flex (Thermo Fisher Scientific, Cat. No. 5400630l) for total RNA isolation. RNA quality was determined using RNA 6000 Nano Assay with BioAnalyzer (Agilent Technologies, Santa Clara, CA), and RNA concentration was determined with Quant-it RNA High Sensitivity assay kit (Thermo Fisher Scientific, Cat. No. Q33140). RNA samples were normalized to 10 ng/ul and stored at-80°C before sequencing. Extracted RNA samples were submitted to the Genome Sequence and Analysis Facility at the University of Texas at Austin for Tag-based RNA sequencing in three separate runs. This method is a cost-effective approach specifically designed to measure the abundances of polyadenylated transcripts, yielding highly reliable data for differential gene expression analysis in well-annotated genomes (Lohman et al., 2016; Meyer et al., 2011). For animals who underwent social hierarchy reorganization, two runs were submitted, one for 70 minutes (N=7/status - descenders, dominant) and another for 25 hours (N=6/status - descenders, dominant). A third run was submitted for animals living in stable social hierarchies (N = 11/status - alpha, subdominant, subordinate). Alpha males refer to males who were rank 1 in each group. Subdominant males are those that had positive David’s Scores, a measure of dominance that indicates that animals tend to win more contests than they lose (Gammell et al., 2003). Subordinate males were either the lowest or second most lowest rank in each vivarium. All runs were processed independently, with libraries constructed with a protocol modified by Meyer et al. (2011) and Lohman et al. (2016). Reads were sequenced on the NovaSeq 6000 SR100 with minimum reads of 4 million and the target reads per sample of 5 million. Raw reads were processed to obtain gene count data by following the TagSeq data processing pipeline based on Meyer et al. (2011) and Lohman et al. (2016). Briefly, a customized Perl script utilizing FASTX-Toolkits and CUTADAPT 2.8 (Martin, 2011) was used to remove reads with a homo-polymer run of “A” ≥ 8 bases and retain reads with a minimum 20 bases and remove PCR duplicates. Processed reads were then mapped to Mus musculus reference genome (Ensembl release 99) using Bowtie2 (Langmead & Salzberg, 2012).

### Experiment 1 – Differential gene expression (DGE) analysis

All DGE analyses were conducted using the Bioconductor package ‘limma’ (Smyth et al., 2021). We first filtered out genes with <10 counts across all samples in each run. Next, filtered read counts were normalized by the trimmed mean of the M-values normalization method (TMM) (Robinson & Oshlack, 2010). Three different DGE analyses were conducted with filtered normalized count data with a voom transform for each batch. For males in the social hierarchy reorganization experiment, DGE analysis identified differentially expressed genes (DEGs) by comparing socially descending alpha males (DES) versus alpha males that remained dominant (DOM) 70 minutes after social reorganization. The same comparison was used with animals sacrificed 25 hours after social reorganization. In the stable social hierarchy experiment, DGE analysis identified DEGs across two comparisons: subordinate (rank 10 - SUB) versus dominant (rank 1 - DOM) males and subdominant (rank 2 or 3 - SUBDOM) versus DOM males. Raw p-values for each DGE analysis were adjusted via empirical false discovery rate (eFDR) (Storey & Tibshirani, 2003). We permuted sample labels 5000 times and obtained a null distribution of p-values to estimate the empirical false discovery rate. We set a threshold for differentially expressed genes as 15% change of the absolute values of log2 fold change at the empirical false discovery rate (eFDR) of 5%. Genes investigated in **Figure 1** and **Supplemental Table 1** represent key genes involved in cholinergic signaling, myelin regulation, and thyroid signaling that were previously identified as top differentially expressed genes in the MeA of descending animals at 70 min (Milewski et al., 2025).

**Figure 1:**
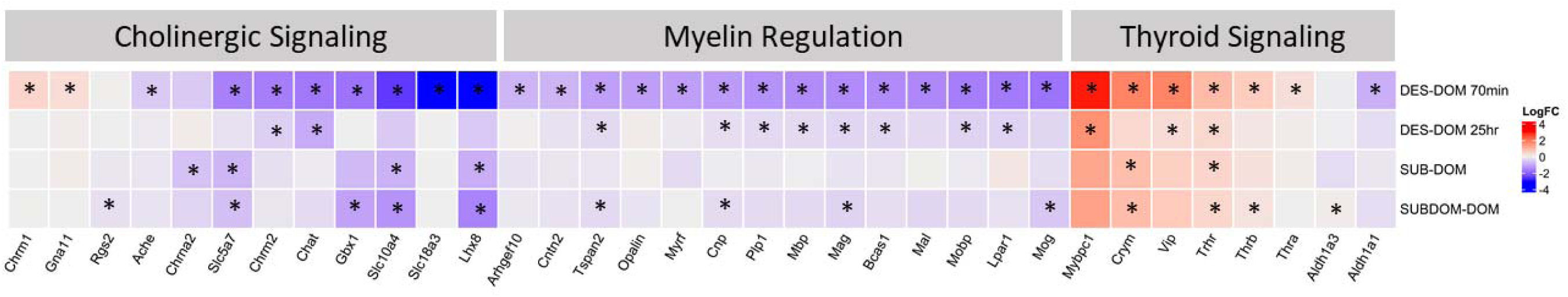
Heatmap of Tag-based RNA sequencing gene expression in reorganized and stable social hierarchies. Relative expression of genes related to cholinergic signaling, myelin regulation, and thyroid signaling are mapped to 4 different comparisons: 1) Descending (DES) to animals who maintained dominance (DOM) at 70 minutes after social reorganization, 2) DES to DOM at 25 hours after a social reorganization, 3) Stable subordinate (SUB) to stable dominant (DOM), and 4) Stable subdominant (SUBDOM) to stable dominant (DOM). Red represents relatively higher gene expression and purple represents relatively lower gene expression in the first group listed in each comparison. Significantly different genes in each comparison with fold changes above 0.2 are represented with a star (*).

### Experiment 2 – Animals, Housing, and Behavioral Observations

In this experiment we utilized RNAscope to assess the spatial distribution of gene expression of specific genes related to cholinergic signaling, myelination, and thyroid signaling that we identified in experiment 1 as differentially expressed between dominant and socially descending males. Male CD-1 mice (N = 144) underwent the identical social hierarchy reorganization paradigm as described in experiment 1. This generated 9 cages of four alpha males who reorganized their social hierarchies. From each cage, 70 minutes following social hierarchy reorganization, the male who maintained dominance status as rank 1 in the cage (DOM) and the male who descended to rank 4 (DES) were sacrificed and whole brains were collected and flash-frozen in a hexane cooling bath, placed on dry ice, and stored at-80 until further processing.

### Experiment 2 – RNAscope *In Situ* Hybridization

Brains were sectioned on a cryostat (Lecia Biosystems, Deer Park, IL) at 20µm at-20, and 2 to 3 sections of the medial amygdala (MeA) were mounted to a slide for animals who stayed socially dominant (DOM, N=9) and for animals who socially descended (DES, N=9) during social reorganization. Slides were stored at-80 until they were processed for RNAscope. RNAscope *in-situ* hybridization (ISH) HiPlex version 2 was performed as instructed by Advance Cell Diagnostics (ACD,(Phatak et al., 2022)). Slides were removed from the-80 freezer and immediately transferred to cold (4°C) 10% formalin for 15 minutes. The tissues were then dehydrated in 50% ethanol (5 min), 70% ethanol (5 min), and 100% ethanol (10 min) at room temperature. The slides were air-dried overnight, and then boundaries were drawn around each section using a hydrophobic pen (ImmEdge PAP pen; Vector Labs). When hydrophobic boundaries had dried, protease IV reagents were added to each section until fully covered and incubated for 30 min at room temperature. Slides were washed with 1X phosphate-buffered saline (PBS) at room temperature as instructed by ACD. All slides were transferred to a prewarmed humidity control tray (ACD) containing dampened paper. Each section was submerged with a mixture of Channel 1 - Fluro: 488, Channel 2 - Dylight: 550, and Channel 3 - Dylight: 650 probes (50:1:1 dilution, as directed by ACD due to stock concentrations) before being incubated in a HybEZ oven (ACD) for 2 hours at probe hybridization. Following probe hybridization, all slides were washed with 1X RNAscope wash buffer (ACD), covered with Hiplex AMP-1 reagent, and returned to the oven for a 30-minute incubation period at 40°C. Washes and amplification were repeated using AMP-2 and AMP-3 reagents with 30-minute incubation periods. After amplification, slides remained in humidity control try, and RNAscope Hiplex Fluoro T1-T3 (*crym*, *mog*, *mbp*) probes were added to cover each section entirely and then slides incubated for 15 mins in the oven at 40°C and then washed with 1X wash buffer at room temperature as instructed by ACD. Slides were stained with DAPI for 30 seconds in the dark and then cover-slipped with Prolong Gold Antifade mounting medium. All sections had left and right sides of 4 subregions: anteroventral medial amygdala (MeAav), anterodorsal medial amygdala (MeAad), basal lateral amygdala BLA, and central amygdala CEA of the amygdala, imaged at 20X using a Leica DMi8 inverted microscope and LASX software (Leica Microsystems). After slides were imaged, they were submerged in 4X SSC (saline-sodium citrate) buffer for 30 to 60 minutes at room temperature. Then, fluorophores were cleaved with freshly prepared 10 % cleaving solution (ACD) at room temperature for 15 minutes and washed in PBST at room temperature. This process was done twice as ACD instructed. The same procedure was repeated for RNAscope Hiplex Fluoro T4-T6 (*mybpc1, fosb, erg3*) and RNAscope HiPlex Fluoro T7-T9 (*chat*, *slc5a7*, *ache*). Target information can be found in **Supplemental Table 2**. Immediate early genes (*fosb* and *erg3*) were stained during RNAscope but were not used in this analysis.

### Experiment 2 – Image Processing and Cell Counting

After all three rounds of RNAscope were completed, the images for each section were overlaid using the DAPI signal as a reference with ACD RNAscope HiPlex Image Registration Software. The resulting 10-channel image (one DAPI channel and a channel of each of the nine probes) was imported into the LASX software for semi-automated quantification. Only high-quality tissue sections absent of wrinkles or damage were counted. As a result, 2 to 3 brain sections were imaged for each subject, leaving 38 (18 left, 20 right) images for both DOM and DES subjects. The LASX Software 2D Analysis module was used to remove the background signal, calculate the total cell count, and quantify the presence of each of the nine probes within each cell for all amygdala subregions. Amygdala subregions were manually outlined using the Allen Brain Atlas for mice as a reference. We used binomial general linear mixed-effect models (GLMMs), including the presence or absence of each target as the outcome variable, social condition (DES vs. DOM) as a fixed effect, and batch, individual ID, and side of the brain as random effects to test the relationship between each target and social condition after social reorganization. GLMMs were created for each target across all four brain subregions of the amygdala. **Table 1** shows the beta coefficients and standard errors for each of the targets of interest. Appropriate GLMMs and LMMs were used for each analysis according to the data distribution and residuals from fitted models using the R package ‘lme4’ (Bates et al., 2023). We used the package ‘lmeRTest’ (Kuznetsova et al., 2020) to derive p-values for GLMMs and LMMs.

**Table 1:**
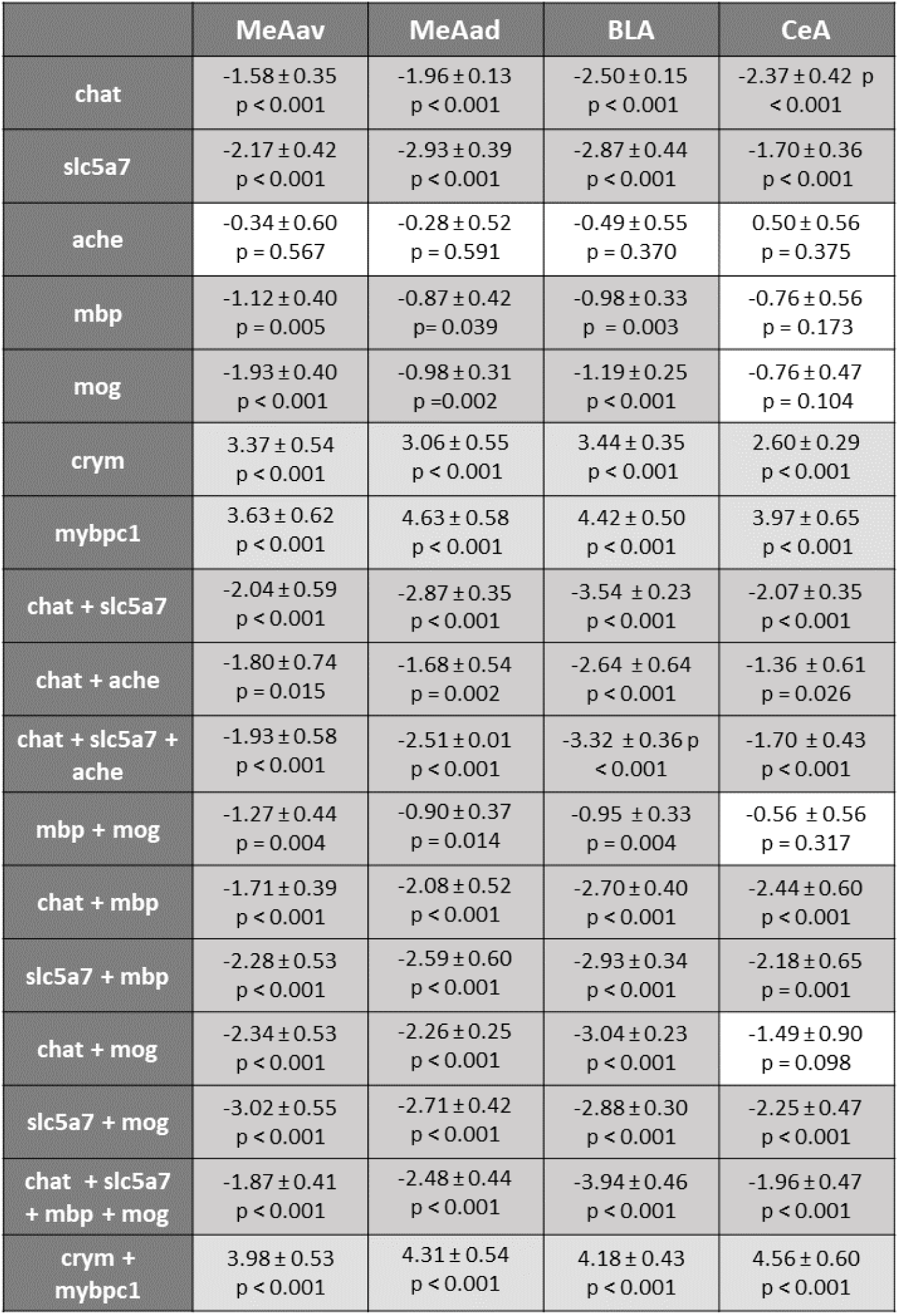
Statistics for all individual and co-expressed mRNA targets. All betas ± standard errors and p-values are reported for each mRNA target across all amygdala subregions. MeAav = anterior ventral medial amygdala, MeAad = anterior dorsal medial amygdala, BLA = basolateral amygdala, and CeA = central amygdala. White = no significant difference, light grey = higher in descenders, and dark grey = higher in dominants.

|  | MeAav | MeAad | BLA | CeA |
| --- | --- | --- | --- | --- |
| chat | -1.58 ± 0.35<br>p < 0.001 | -1.96 ± 0.13<br>p < 0.001 | -2.50 ± 0.15<br>p < 0.001 | -2.37 ± 0.42 p<br>< 0.001 |
| slc5a7 | -2.17 ± 0.42<br>p < 0.001 | -2.93 ± 0.39<br>p < 0.001 | -2.87 ± 0.44<br>p < 0.001 | -1.70 ± 0.36<br>p < 0.001 |
| ache | -0.34 ± 0.60<br>p = 0.567 | -0.28 ± 0.52<br>p = 0.591 | -0.49 ± 0.55<br>p = 0.370 | 0.50 ± 0.56<br>p = 0.375 |
| mbp | -1.12 ± 0.40<br>p = 0.005 | -0.87 ± 0.42<br>p = 0.039 | -0.98 ± 0.33<br>p = 0.003 | -0.76 ± 0.56<br>p = 0.173 |
| mog | -1.93 ± 0.40<br>p < 0.001 | -0.98 ± 0.31<br>p = 0.002 | -1.19 ± 0.25<br>p < 0.001 | -0.76 ± 0.47<br>p = 0.104 |
| crym | 3.37 ± 0.54<br>p < 0.001 | 3.06 ± 0.55<br>p < 0.001 | 3.44 ± 0.35<br>p < 0.001 | 2.60 ± 0.29<br>p < 0.001 |
| mybpc1 | 3.63 ± 0.62<br>p < 0.001 | 4.63 ± 0.58<br>p < 0.001 | 4.42 ± 0.50<br>p < 0.001 | 3.97 ± 0.65<br>p < 0.001 |
| chat + slc5a7 | -2.04 ± 0.59<br>p < 0.001 | -2.87 ± 0.35<br>p < 0.001 | -3.54 ± 0.23<br>p < 0.001 | -2.07 ± 0.35<br>p < 0.001 |
| chat + ache | -1.80 ± 0.74<br>p = 0.015 | -1.68 ± 0.54<br>p = 0.002 | -2.64 ± 0.64<br>p < 0.001 | -1.36 ± 0.61<br>p = 0.026 |
| chat + slc5a7 +<br>ache | -1.93 ± 0.58<br>p < 0.001 | -2.51 ± 0.01<br>p < 0.001 | -3.32 ± 0.36 p<br>< 0.001 | -1.70 ± 0.43<br>p < 0.001 |
| mbp + mog | -1.27 ± 0.44<br>p = 0.004 | -0.90 ± 0.37<br>p = 0.014 | -0.95 ± 0.33<br>p = 0.004 | -0.56 ± 0.56<br>p = 0.317 |
| chat + mbp | -1.71 ± 0.39<br>p < 0.001 | -2.08 ± 0.52<br>p < 0.001 | -2.70 ± 0.40<br>p < 0.001 | -2.44 ± 0.60<br>p < 0.001 |
| slc5a7 + mbp | -2.28 ± 0.53<br>p < 0.001 | -2.59 ± 0.60<br>p < 0.001 | -2.93 ± 0.34<br>p < 0.001 | -2.18 ± 0.65<br>p = 0.001 |
| chat + mog | -2.34 ± 0.53<br>p < 0.001 | -2.26 ± 0.25<br>p < 0.001 | -3.04 ± 0.23<br>p < 0.001 | -1.49 ± 0.90<br>p = 0.098 |
| slc5a7 + mog | -3.02 ± 0.55<br>p < 0.001 | -2.71 ± 0.42<br>p < 0.001 | -2.88 ± 0.30<br>p < 0.001 | -2.25 ± 0.47<br>p < 0.001 |
| chat + slc5a7<br>+ mbp + mog | -1.87 ± 0.41<br>p < 0.001 | -2.48 ± 0.44<br>p < 0.001 | -3.94 ± 0.46<br>p < 0.001 | -1.96 ± 0.47<br>p < 0.001 |
| crym +<br>mybpc1 | 3.98 ± 0.53<br>p < 0.001 | 4.31 ± 0.54<br>p < 0.001 | 4.18 ± 0.43<br>p < 0.001 | 4.56 ± 0.60<br>p < 0.001 |

### Experiment 2 – Social Behavior Analysis

Win/loss sociomatrices for each social group were created pre-and post-reorganization using the frequency of wins and losses observed from behavioral observations (**Supplemental Figure 2**). From these sociomatrices, we tested the stability of each social hierarchy by testing the significance of directional consistency as previously described (Williamson et al., 2016; Williamson et al., 2019). Individual social ranks were determined by David’s Scores (DS). DS is a measure of individual dominance related to the proportion of wins to losses adjusted for opponents’ relative dominance (Gammell et al., 2003). DS was calculated pre-reorganization and at 70 min after reorganization using the R package ‘compete’ (Curley, 2016). We used linear mixed-effect models (LMMs) to determine if body weight was related to DS before reorganization. Each LMM had pre-reorganization DS as the outcome variable, with the fixed effect being each individual’s body weight at GD1 or GD11. We also included individual ID, batch, and cage ID as random effects. Additional LMMs were used to determine which variables were predicted DS after reorganization. We also included a fixed effect, ‘total aggression rate per cage’, which was the rate per hour of all aggression in each individual’s cage before the reorganization. This was to test if behavioral dynamics within the cage-of-origin influenced post-reorganization DS. Each LMM had post-reorganization DS as the outcome variable, with the fixed effect being each individual’s body weight at GD1, GD11, or rate of aggression given or received by each mouse in their pre-reorganization cage. Rates of aggression given were calculated as each individual’s total number of aggressive acts given or received per hour. Random effects were individual ID, batch, and cage ID.

## Results

### Experiment 1 – Tag-based RNA Sequencing in stable and reorganized hierarchies

#### Expression of cholingeric signaling genes is relatively higher in dominant males in stable and reorganized social hierarchies

Previously, we found that 7 out of the top 25 most differentially expressed genes in the DES vs DOM comparison at 70 minutes were acetylcholinergic pathway-related genes (Milewski et al., 2025). Here, we examined whether genes related to this signaling pathway were i) similarly differentially expressed in the MeA 25 hours following a social hierarchy reorganization and ii) were differentially expressed between dominant and males of lower social rank from stable social hierarchies. We found that 12 cholinergic signaling genes showed correlated log-fold changes in dominant males across gene sets (**Figure 1 and Supplemental Table 1**). Socially reorganized dominant males that maintained dominance status showed significantly higher expression in eight of these genes at 70 minutes: LIM homeobox protein 8 (*lhx8*), solute carrier family 18 member A3 (*slc18a3*), solute carrier family 10 member 4 (*slc10a4*), solute carrier family 5 member 7 (*slc5a7*), gastrulation brain homeobox 1 (*gbx1*), choline o-acetyltransferase (*chat*), cholinergic receptor muscarinic 2 (*chrm2*), and acetylcholinesterase (*ache*). Only two (*chrm2*, *chat*) of these genes continued to have significantly higher expression in DOM males compared to DES males 25 hours after social reorganization (**Figure 1 and Supplemental Table 1**). *chrm1* (cholinergic receptor muscarinic 1) and *gna11* (G protein subunit alpha 11) exhibited relatively lower expression in dominant males 70 minutes after reorganization but were insignificant in all other comparisons. Two solute carriers (*slc10a4* and *slc5a7*) and *lhx8* were more highly expressed in stable dominant males than in both subdominant and subordinate males. Additionally, *chrna2* (cholinergic receptor nicotinic alpha 2 subunit) was significantly higher in dominant males compared to subordinate males in stable hierarchies. These data suggest that the upregulation of cholinergic functioning may underlie both the rapid acquisition and maintenance of dominance phenotypes.

#### Reduced expression of genes associated with myelination regulation is specifically associated with social descent

Previously, we observed elevated expression in many myelin-related genes in the MeA of alpha males who maintained dominant status 70 minutes after social reorganization compared to those who were socially descended (Milewski et al., 2025). Of the 14 myelin-related genes found to be higher in dominants 70 minutes after reorganization, eight (*tspan2* - Tetraspanin 2, *cnp* - 2’,3’-Cyclic Nucleotide 3’ Phosphodiesterase, *plp1* - proteolipid protein 1, *mbp* - myelin basic protein, *mag* - myelin-associated glycoprotein, *bcas1* - brain enriched myelin associated protein 1, *mobp* - myelin-associated oligodendrocyte basic protein, *lpar1* - lysophosphatidic acid receptor 1) still had significantly higher expression in dominant animals 25 hours after social reorganization **(Figure 1 and Supplemental Table 1)**. None of the myelination genes were upregulated in stable dominants compared to subordinate males, although four (*tspan2*, *cnp*, *mag*, *mog* - myelin oligodendrocyte glycoprotein) were over-expressed in stable dominants compared to subdominant males. These gene expression patterns suggest that upregulation of myelination is essential for maintaining dominance during social challenges but not necessarily in stable social hierarchies.

#### Expression of thyroid signaling associated genes is reduced in dominant males

We examined eight thyroid-signaling associated genes in the MeA across socially disrupted and stable datasets **(Figure 1 and Supplemental Table 1)**. All genes exhibited consistent patterns of fold changes across comparisons. Six genes had relatively higher expression in socially descending males and subordinate males compared to dominant males. Myosin binding protein C1 (*mybpc1*) and vasoactive intestinal peptide (*vip*) were significantly higher in descending animals at both 70 minutes and 25 hours but showed no significant differences in stable animals. Crystallin Mu (*crym)* was significantly more highly expressed in descending males at 70 minutes but not 25 hours after social reorganization. This gene was also significantly more highly expressed in subdominant and subordinate males compared to dominant males in stable hierarchies. Both thyroid hormone receptor alpha (*thra*) and beta (*thrb*) were significantly higher in descending animals at 70 minutes, and *thrb* was also significantly more highly expressed in descending males at 25 hours post-reorganization. Both of these genes also show significantly higher expression in subordinates compared to dominant males living in stable hierarchies. Notably, *trhr* (thyrotropin-releasing hormone receptor gene) was significantly more highly expressed in descending animals than dominant animals at both time-points as well as being significantly higher in stable subordinate and subdominant males compared to stable dominant males **(Figure 1 and Supplemental Table 1)**. Together, these gene expression patterns suggest that genes associated with the thyroid signaling pathway are activated in more subordinate animals and genes engaged in this pathway increase expression as animals start to fall in social status.

### Experiment 2 – Assessing gene expression in the amygdala using *in situ* hybridization in males who socially descend versus maintain dominance

#### Mice rapidly reform social hierarchies following a social reorganization

Before groups were socially reorganized, all 36 cages of four male mice formed stable linear hierarchies with highly significant directional consistency with more dominant animals directing aggression toward more subordinate animals (**Supplemental Figure 2**, median[IQR] = 0.91[0.86,0.96], all p <0.001). Clear individual ranks were observed in each cage’s hierarchy based on David’s score (**Supplemental Figure 3A**). Alpha males (rank 1), had the highest positive David’s score (median[IQR] = 4.99[4.50,5.40]). Beta males (rank 2) had David’s Scores that were mostly positive (median[IQR] = 0.41[-0.03,1.10]), indicating that, on average, beta males win more contests than they lose. Gamma males (rank 3, median[IQR] =-1.80[-2.18,-1.43]) and delta males (rank 4, median[IQR] =-3.51[-4.32,-2.52]) all had David’s scores below zero, demonstrating that they lost more contests than they won. Body mass on day 1 (**Supplemental Figure 3B**, β=0.28±0.11, p=0.012) and day 11 (**Supplemental Figure 3C**, β=0.35±0.10, p < 0.001) of group-housing was significantly moderately associated with David’s score. On GD11, alpha males weighed significantly more than beta (β=-1.38±0.53, p=0.011), gamma (β=-1.31±0.53, p=0.015), and delta (β=-1.40±0.53, p =0.009) males. There were no significant differences between other social ranks in body mass.

After alpha males were placed into a new cage of four males, they rapidly reformed social hierarchies within 70 minutes of social reorganization with highly significant directional consistency (**Supplemental Figure 4A**, median[IQR] = 0.75[0.68,0.78], p <0.001). Although all animals were previously dominant, new social ranks emerged as groups were reorganized. Some males remained dominant with positive David’s scores (**Supplemental Figure 4A**, median[IQR] = 4.01[3.86,4.94]) whereas others socially descended to rank 2 (median[IQR] = 0.77(-0.15,1.89]), rank 3 (median[IQR] =-1.55[-1.75,-1.17]), or rank 4 (median[IQR] =-3.91[-4.25,-2.95]). Neither body mas on GD1 or GD11 nor David’s score in the preorganized cage significantly predicted an individual’s David’s score in the new hierarchy (**Supplemental Figure 4B & 4C**). We also found that individual aggression given in the pre-reorganization cage did not predict post-reorganized David’s Score although there was a trend or eventual alpha males to have lower aggression levels (**Supplemental Figure 4D**). However, we did find that total cage aggression in an individual’s cage before social reorganization did predict David’s Score 70 minutes after reorganization with alpha males from less aggressive cages being more likely to establish dominance in their new hierarchy (β=-0.07±0.03, p=0.032; r(CI) =-0.39(-0.64,-0.07), p = 0.019; **Supplemental Figure 4E**).

#### Social descent is associated with specific patterns of gene expression in distinct amygdaloid subnuclei

We employed RNAscope to map mRNA markers of social descent versus dominance across four amygdala subregions. We used RNA markers *chat*, *slc5a7*, and *ache* to evaluate the cholinergic signaling pathway. This allowed for investigation of acetylcholine’s formation, movement, and dispersal within neurons. The RNA markers used for myelination were *mbp* and *mog* to identify mature oligodendrocytes. Finally, *crym* was used as the RNA marker to examine thyroid signaling, as it regulates the amount of freely available thyroid hormone triiodothyronine (T3) (Aksoy et al., 2022). All of these genes have previously been determined to be some of the most differentially expressed between dominant and descending males using Tag-based RNA sequencing (Milewski et al., 2025). Similarly, we used *mybpc1* as an RNA target as it was the most highly expressed DEG in descending males. *mybpc1* is not known to have a direct connection to the thyroid signaling pathway but is a marker for vasoactive intestinal polypeptide-expressing (*vip*) interneurons in the cortex (Jiang et al., 2023), and *vip* gene expression is also regulated by thyroxine (T4) hormone in the pituitary (Segerson et al., 1989), suggesting that the thyroid signaling pathway may regulate *mybpc1* too.

**Table 1** shows statistics for all individual targets and co-expressed targets.

Representative images are shown in **Figure 2** and **Supplemental Figure 5**. Animals who maintained their dominance during social reorganization exhibited a significantly higher proportion of cells expressing *chat* and *slc5a7* across all four medial amygdala regions (**Figure 3**). A significantly higher proportion of cells co-expressed both *chat+slc5a7* in the MeAav, MeAad, BLA, and CeA in dominant animals compared to descenders (**Figure 4**). In contrast to our Tag-based RNA sequencing data, we did not find that dominant animals express a higher proportion of *ache* positive cells than descending males in any of the amygdala subregions.

**Figure 2.**
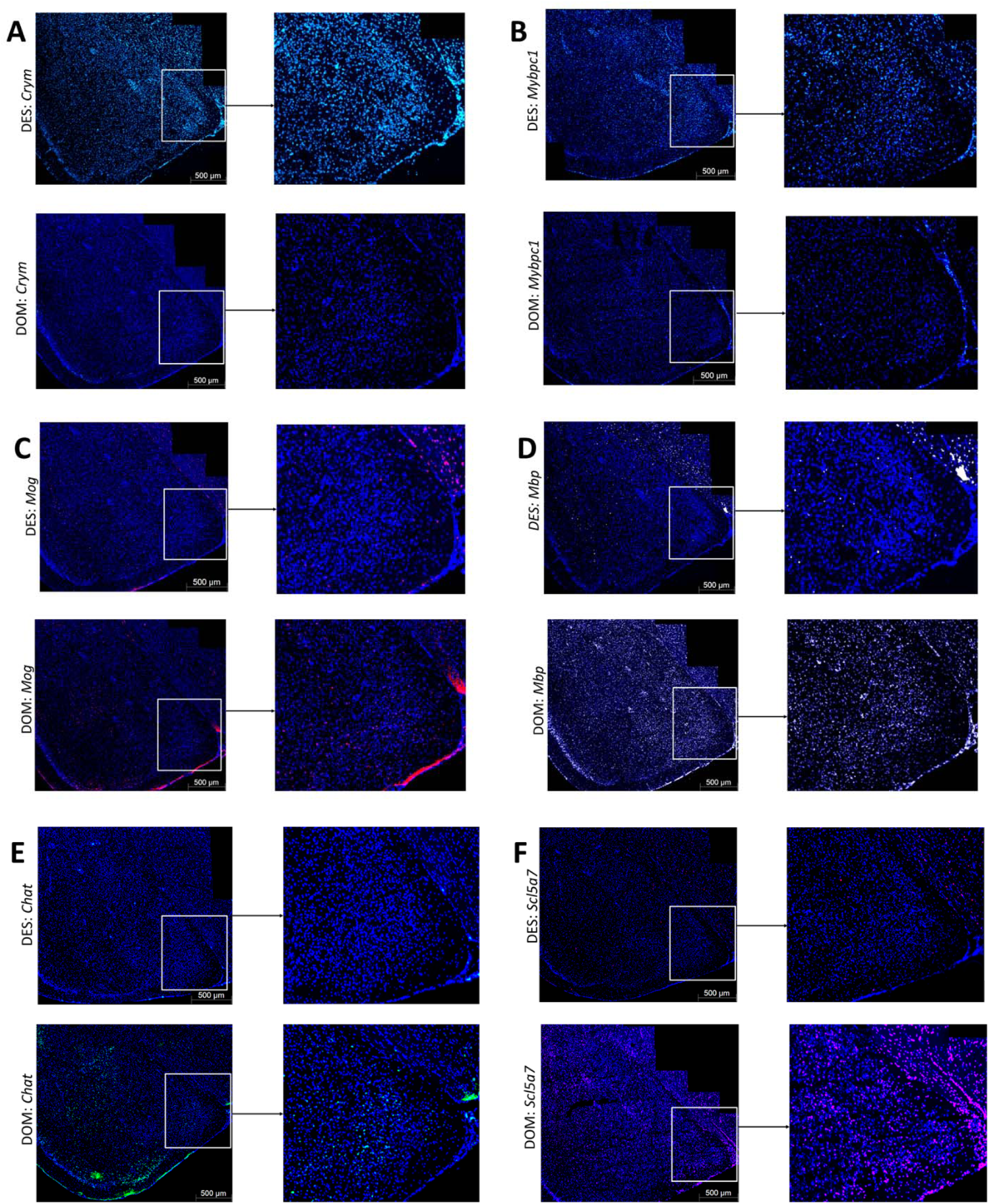
Representative images of mRNA expression of **A)** *crym* **B)** *mybpc1*, **C)** *mog*, **D)** *mbp,* **E)** *chat*, **F)** *slc5a7* in the anterior MeA in descending (DES) versus dominant (DOM) males.

**Figure 3:**
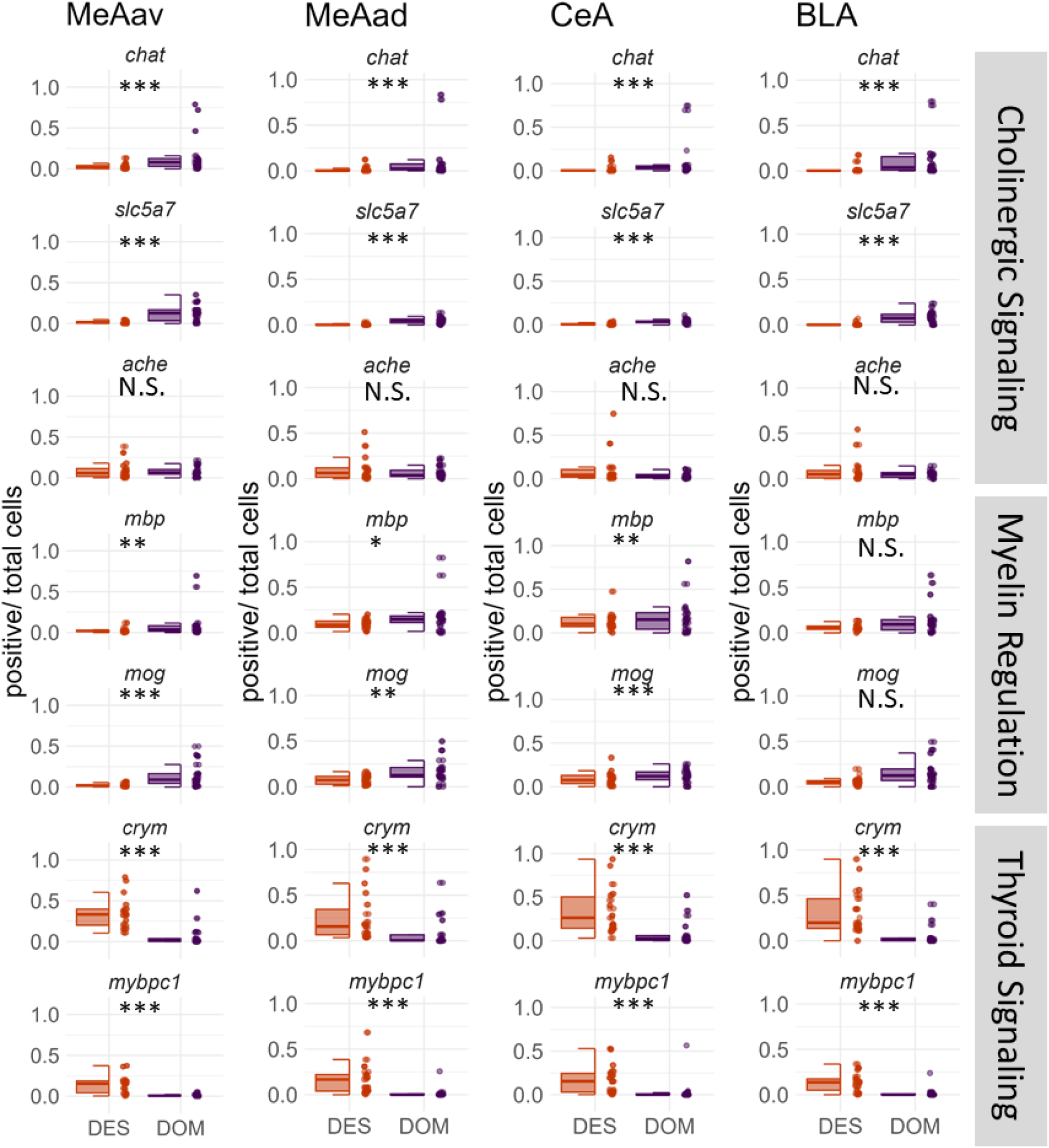
Expression of select genes in subnuclei of the amygdala using RNAscope. Boxplots represent the proportion of activated cells for each mRNA target from total cells by social condition (descending males versus dominant males) for each amygdala subregion. Points represent individual sections. DES = dominant animals who descended, DOM = animals who maintained dominance, MeAav = anterior ventral medial amygdala, MeAad = anterior dorsal medial amygdala, BLA = basolateral amygdala, CeA = central amygdala. * p<0.05, ** p<0.001, *** p<0.001

**Figure 4:**
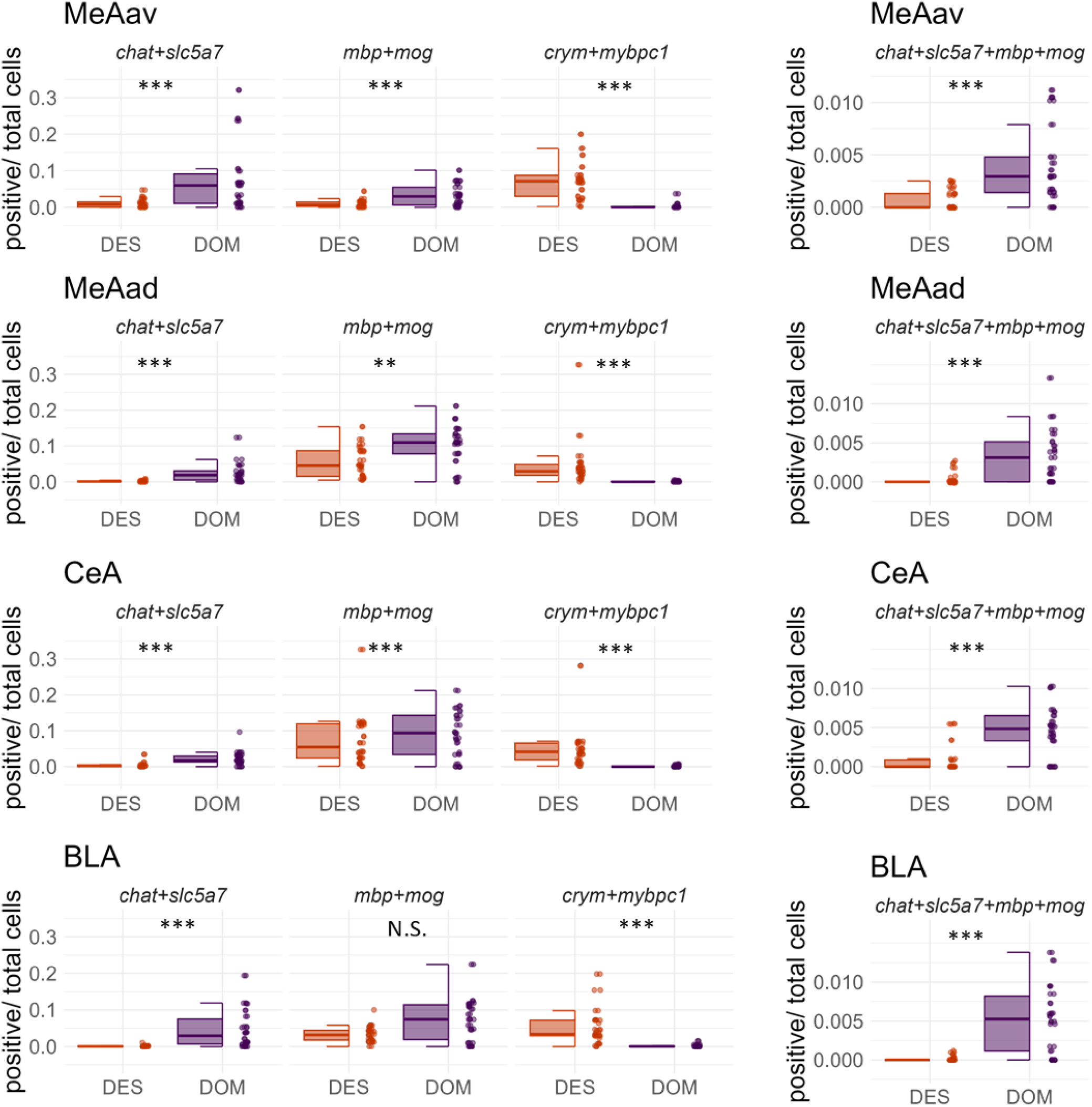
Co-expression of mRNA targets from RNAscope. Boxplots represent the proportion of cells showing co-expression from total cells by social condition (descending males versus dominant males) for each amygdala subregion. Points represent individual subjects. DES = dominant animals who descended, DOM = animals who maintained dominance, MeAav = anterior ventral medial amygdala, MeAad = anterior dorsal medial amygdala, BLA = basolateral amygdala, CeA = central amygdala. * p<0.05, ** p<0.001, *** p<0.001

However, dominant males did have a significantly higher proportion of cells co-expressing *chat+ache,* as well as all three genes *chat+slc5a7+ache* in the MeA, BLA, and CeA (**Table 1**).

Dominant males also had significantly higher proportion of cells expressing *mog* and *mbp* in the anterior MeA and BLA compared to descending males (**Figure 3**, **Table 1**) as well as a higher proportion of cells co-expressing *mog*+*mbp* in these regions (**Figure 4**). There were no significant differences in these markers in the CeA. In all four regions there was a large significant reduction in the proportion of cells co-expressing *chat+mbp*, *slc5a7+mbp* or *slc5a7+mog* in descending males (**Table 1**). In the anterior MeA and BLA there were also large significant reductions in the proportion of cells co-expressing *chat*+*mog* in descending males.

We found extremely large differences between dominant and descending males in the expression of *crym* and *mybpc1* in all four subnuclei of the amygdala. Descending males had significantly elevated expression of both markers as well as a significantly higher proportion of cells co-expressing *crym+mybpc1* compared to dominant males (**Figures 3 & 4**, **Table 1**).

## Discussion

We previously identified three pathways (cholinergic signaling, myelin regulation, thyroid hormone signaling) that rapidly alter their levels of gene expression in the MeA 70 minutes following animals changing social status. Here, we further investigate these pathways by characterizing how the expression of these genes is modulated in the MeA 25 hours following social rank change. Further, we examined the expression of genes related to these pathways in dominant and subordinate male mice living in stable social hierarchies to determine if the observed gene expression patterns are specifically associated with changing status or are more generally associated with individual ranks. We mapped the expression of 12 genes related to cholinergic signaling, 14 genes involved in myelin regulation, and 8 genes affiliated with thyroid signaling from MeA Tag-based RNA sequencing datasets. Further, we use RNAscope, an *in-situ* hybridization protocol that allows for multiple targets, to map 7 of these gene targets (*chat, slc5a7, ache, mbp, mog, crym, mybpc1*) and their co-expression spatially in four subregions of the amygdala.

Our data suggest that elevated cholinergic functioning may help support and maintain a dominant phenotype during social reorganization and stable social hierarchies. Cholinergic signaling is highly activated 70 minutes after social reorganization. Animals that retained dominant status showed elevated expression of *chat*, the gene encoding choline acetyltransferase which is responsible for the synthesis of acetylcholine in neurons. They also exhibited higher expression of vesicular acetylcholine transporters (*slc18a3*, *slc5a7*, *slc10a4*), which are essential membrane proteins that package and transport acetylcholine into synaptic vesicles in cholinergic neurons. Additionally, increased expression in males that maintain dominance was observed for the muscarinic receptor *chrm2* and *ache*, the gene responsible for producing acetylcholinesterase, the enzyme that rapidly breaks down acetylcholine in the synaptic cleft. Dominant animals also showed elevated expression of the homeobox gene *lhx8*, which plays a key role in the development of cholinergic neurons (Asbreuk et al., 2002; Mori et al., 2004) and promotes vesicular acetylcholine transporter function and acetylcholine release (Li et al., 2014; Tomioka et al., 2014). *Chat* and *chrm2* are still elevated in animals who maintained dominance compared to descending males 25 hours after social reorganization. RNAscope results confirmed *chat* and *slc5a7* activation in the MeA of male mice who maintained dominance 70 minutes after social reorganization. However, unlike the sequencing data, we found no significant activation with *ache* alone. We did see the co-expression of *chat+slc5a7+ache* in a significantly higher proportion of cells of male mice maintaining dominance though this effect may be largely driven by the higher expression of *chat* and *slc5a7*. Importantly, the activation of *chat, slc5a7,* and the co-expression of all three cholinergic mRNA targets was not limited to the MeA but also detected in the BLA and CeA subnuclei of the amygdala. In stable hierarchies, dominant males have higher expression of cholinergic vesicle carriers *slc10a4* and *slc5a7* as well as *lhx8,* which modulates the activity of these vesicle carriers (Li et al., 2014; Tomioka et al., 2014) compared to stable subordinate and subdominant males. Taken together, these data suggest that high levels of cholinergic signaling may be required to maintain ongoing dominance in both stable and reorganized hierarchies.

Cholinergic fibers innervate throughout subnuclei of the amygdala and activate both glutamatergic and GABAergic neurons, consequently influencing the amygdala’s inhibitory/excitatory balance (Jie et al., 2018). Cholinergic functioning in the forebrain and amygdala modulates several social behaviors, including social stress responsivity (Mineur et al., 2016), social recognition (Prado et al., 2006), social memory formation (Kljakic et al., 2021; Winslow & Camacho, 1995), and fear learning (Crimmins et al., 2023). Of these, probably the most well characterized is its role in the formation and extinction of fear memories in the BLA (Crimmins et al., 2023; Kellis et al., 2020). *Chat+* neurons from the forebrain innervate the BLA-releasing acetylcholine (Aitta-aho et al., 2018), and update their activity when the valence of stimuli changes (Saddoris et al., 2005). *chat+* neurons from the brainstem also innervate the CeA and induce defensive learning-related behaviors (Aitta-aho et al., 2018). Functional roles of vesicle carriers are less well-established (Ribeiro et al., 2006), but *slc5a7* mediates choline uptake at the presynaptic neuron terminal and likely influences cholinergic signaling. Although less is known regarding cholinergic innervation in the MeA compared to other amygdala nuclei (Hecker & Mesulam, 1994), our findings suggest cholinergic activity in the MeA and other sub nuclei of the amygdala may contribute to the success of dominant animals maintaining their dominance status during social challenges.

Reduced expression of genes associated with myelination was consistently associated with animals rapidly descending in social status during social reorganization. In contrast, myelin activation appears largely unrelated to the ongoing maintenance of dominance in stable hierarchies. Differential expression analysis revealed 14 myelin-related genes were upregulated in males who maintained their dominant status compared to those who descended at 70 minutes. Eight of these genes continued to be differentially expressed at 25 hours post social reorganization, suggesting that changes in myelination are persistent even after the new hierarchy has been established. These genes include those that affect myelin structure and composition, such as *mbp*, *mobp*, and *mag*; genes involved in the development and regulation of myelin and oligodendrocytes (*mog*, *mal*); and others associated with myelin-related signaling and cellular interactions, including *lapr1* (lysophosphatidic acid receptor 1), *tspan2* (tetraspanin-2), *cntn2* (contactin 2), and *arhgef10* (rho guanine nucleotide exchange factor 10). Several studies have demonstrated that chronic social stress can lead to decreased expression of multiple myelin-associated genes, resulting in long-term reductions in oligodendrocyte populations, as well as diminished levels and thickness of myelin proteins in regions such as the mPFC, hippocampus, and nucleus accumbens (Bonnefil et al., 2019; Lehmann et al., 2017). Social stressors have also been shown to reduce the expression of oligodendrocyte genes encoding myelin proteins in the BLA and CeA (Cathomas et al., 2019). Chronic social stress may also increase levels of myelination in a brain region specific manner. For example, stress decreases the density of proliferative oligodendrocyte precursor cells in the mPFC and BLA but increases the density of mature oligodendrocytes in the BLA (Poggi et al., 2022). Notably, the social reorganization in our study is a much shorter social challenge than chronic social stress that typically lasts up to two weeks, yet we identify changes in myelin-related gene expression that occur immediately following the social disruption. Indeed, our RNAscope data confirmed the higher expression of *mbp* and *mog* in the MeA of males who maintained dominant status 70 minutes after social reorganization and identified the same pattern in the BLA but not the CeA. *mbp* and *mog* are markers of mature oligodendrocytes that have distinct roles in the formation and development of myelin (Kuhn et al., 2019; Sams, 2021). Recently, there has been increased attention to the processes through which oligodendrocytes modulate neuronal function beyond altering axonal conductance, such as by influencing synaptic efficacy, excitability, and plasticity (Xin & Chan, 2020). Indeed, it has been proposed that altered myelination patterns provide a mechanism through which experience-based learning can modify existing neural circuits (Bonetto et al., 2021). It is, therefore, possible that the rapid changes in myelination gene expression we observe in socially descending animals are required for these animals to modify neural circuits to adjust to their new social niche.

To determine if there was any overlap in the activity of genes from cholinergic and myelination pathways following social reorganization, we compared the co-expression of genes across cells with RNAscope. We found a significant increase in the proportion of cells expressing both cholinergic and myelin genes (*chat, slc5a7, mbp, and mog)* in males who maintained dominance 70 minutes after social reorganization compared to those who descend. Myelinating glia express muscarinic and nicotinic acetylcholine receptors, and acetylcholinesterase is highly expressed in white matter (Fields, 2015; Fields et al., 2017). It has been proposed that cholinergic activity can therefore lead to increased myelination of neurons (De Angelis et al., 2012; Fields et al., 2017). It is possible that the higher levels of cholinergic signaling we observe in dominant males may promote myelination, however, since we did not use a cell type marker, it remains unclear if these cells are glia. Future work will need to determine if social hierarchy reorganization leads to long-term structural changes in myelin or if these effects on gene expression are transient, and whether these are driven by altered acetylcholinergic tone.

Genes associated with thyroid signaling were found to be relatively more highly expressed in descending males compared to those who maintained their dominant status. RNAscope confirmed that descending males had a significantly higher proportion of cells expressing *crym* and *mybpc1* in the MeA as well as in the BLA and CeA. This effect was dramatic with very little expression of these genes in animals that maintained their dominance. We also observed that subdominant males in stable groups had relatively higher expression of thyroid hormone receptors alpha (*thra*) and beta (*thrb*) as well as *crym* than dominant males in the MeA. In contrast, subordinates had higher expression of both *thra* and *crym* compared to stable dominant males. Thyroid hormones are known to induce learning-associated neuroplasticity in a number of brain regions (Raymaekers & Darras, 2017). Thyroid hormone signaling in the amygdala has been shown to play an important role in facilitating fear learning (Bárez-López et al., 2017; Montero-Pedrazuela et al., 2011), and promoting social vigilance specifically in the MeA (Kwon et al., 2021). Global knockout of thyroid receptors alpha and beta increases and decreases anxiety, respectively in mice (Vasudevan et al., 2013). Further, thyrotropin-releasing hormone is also down-regulated in the BLA of mice exhibiting stress-resiliency compared to those that develop stress-induced depressive-like behaviors (Choi et al., 2005). *Crym* acts as a binding protein for T3 and, therefore, modulates the amount of freely available T3 in the cytosol (Aksoy et al., 2022). Within the MeA, overexpression of *crym* has been shown to coordinate the response to social isolation stress (Walker et al., 2022). Less is known about the function role of *mybpc1* in the amygdala, but it appears to be a molecular marker for a subpopulation of *vip* interneurons in the cortex (Jiang et al., 2023). *Vip* expression is regulated by thyroid hormones (Giardino et al., 1994; Segerson et al., 1989) and is an important regulator of social behaviors including aggression (Kingsbury & Wilson, 2016) and stress coping during social defeat (Mineur et al., 2022). It is possible that thyroid hormone signaling may also influence *vip* and *mybpc1* amygdala expression following social hierarchy reorganization.

Congruent with this, we observed very high co-expression of *crym* and *mybpc1* across all four subregions of the amygdala in descending males. Overall, our findings suggest that dominant males in stable conditions and those facing a social challenge such as a hierarchy reorganization maintain relatively low levels of thyroid hormone signaling in these amygdala subnuclei. We hypothesize that these reduced levels of thyroid hormone signaling help maintain their resilience to social stress, whereas subordinate individuals increase thyroid signaling to facilitate socio-emotional learning related to their new social status.

## Conclusion

In this study, we demonstrate that male mice are able to rapidly reform social hierarchies immediately following social reorganization. We characterize changes in gene expression in cholinergic signaling, myelination, and thyroid signaling pathways in four subnuclei of the amygdala that are distinctly associated with maintaining or changing social status. Using Tag-based RNA sequencing and RNAscope, we identify that cholinergic signaling is highly activated as animals reestablish dominance status during a social reorganization as well as while maintaining dominance status in stable conditions. We also observe higher levels of myelination-related genes in dominant males compared to descenders in the MeA and BLA. High levels of cholinergic activity may initially promote myelination to promote social learning and resilience following this acute social stress. Descending males continue to show consistently reduced levels of myelination genes 25 hours post reorganization while fewer cholinergic signaling genes exhibit reduced expression at this time point. Further, minimal differences in myelin-related genes are observed between dominant and subordinate males in stable hierarchies, suggesting that differences in myelination between ranks are specific to social status changes. Lastly, we find evidence that the expression of genes associated with thyroid signaling is elevated in descending and subordinate males compared to dominant males in both stable and reorganized hierarchies, which may support the social learning required for their adaptation to lower ranks. In summary, rapid modifications in each of these three pathways help shape the social behaviors and responses to social stress required by dominant and subordinate mice to facilitate their adaptations to changing social environments.

## Data Availability

Data and R code used in the analyses of this paper are available at: https://github.com/ty14/SocialTransitions_RNAscope

## Notes

https://github.com/ty14/SocialTransitions_RNAscope

### Competing Interest Statement

The authors have declared no competing interest.

## References

Aitta-aho, T., Hay, Y. A., Phillips, B. U., Saksida, L. M., Bussey, T. J., Paulsen, O., & Apergis-Schoute, J. (2018). Basal Forebrain and Brainstem Cholinergic Neurons Differentially Impact Amygdala Circuits and Learning-Related Behavior. Current Biology, 28(16), 2557–2569.e4. 10.1016/j.cub.2018.06.064

Aksoy, O., Hantusch, B., & Kenner, L. (2022). Emerging role of T3-binding protein μ-crystallin (CRYM) in health and disease. Trends in Endocrinology & Metabolism, 33(12), 804–816. 10.1016/j.tem.2022.09.003

Anderson, D. J. (2016). Circuit modules linking internal states and social behaviour in flies and mice. Nature Reviews Neuroscience, 17(11), Article 11. 10.1038/nrn.2016.125

Asbreuk, C. H. J., Vogelaar, C. F., Hellemons, A., Smidt, M. P., & Burbach, J. P. H. (2002). CNS Expression Pattern of Lmx1b and Coexpression with Ptx Genes Suggest Functional Cooperativity in the Development of Forebrain Motor Control Systems. Molecular and Cellular Neuroscience, 21(3), 410–420. 10.1006/mcne.2002.1182

Balázs, R., Brooksbank, B. W. L., Davison, A. N., Eayrs, J. T., & Wilson, D. A. (1969). The effect of neonatal thyroidectomy on myelination in the rat brain. Brain Research, 15(1), 219–232. 10.1016/0006-8993(69)90321-7

Bárez-López, S., Montero-Pedrazuela, A., Bosch-García, D., Venero, C., & Guadaño-Ferraz, A. (2017). Increased anxiety and fear memory in adult mice lacking type 2 deiodinase. Psychoneuroendocrinology, 84, 51–60. 10.1016/j.psyneuen.2017.06.013

Bates, D., Maechler, M., Bolker, B., Walker, S., Christensen, R. H. B., Singmann, H., Dai, B., Scheipl, F., Grothendieck, G., Green, P., Fox, J., Bauer, A., & Pavel, K. (2023). lme4: Linear Mixed-Effects Models using “Eigen” and S4 (Version 1.1-34) [Computer software]. https://cran.r-project.org/web/packages/lme4/index.html

Bell, R., Warburton, D. M., & Brown, K. (2009). Drugs as Research Tools in Psychology: Cholinergic Drugs and Aggression. Neuropsychobiology, 14(4), 181–192. 10.1159/000118225

Bonetto, G., Káradóttir, R., & Belin, D. (2021). Myelin: A gatekeeper of activity-dependent circuit plasticity? Science, 374. 10.1126/science.aba6905

Bonnefil, V., Dietz, K., Amatruda, M., Wentling, M., Aubry, A. V., Dupree, J. L., Temple, G., Park, H.-J., Burghardt, N. S., Casaccia, P., & Liu, J. (2019). Region-specific myelin differences define behavioral consequences of chronic social defeat stress in mice. eLife, 8, e40855. 10.7554/eLife.40855

Cathomas, F., Azzinnari, D., Bergamini, G., Sigrist, H., Buerge, M., Hoop, V., Wicki, B., Goetze, L., Soares, S., Kukelova, D., Seifritz, E., Goebbels, S., Nave, K.-A., Ghandour, M. S., Seoighe, C., Hildebrandt, T., Leparc, G., Klein, H., Stupka, E.,… Pryce, C. R. (2019). Oligodendrocyte gene expression is reduced by and influences effects of chronic social stress in mice. Genes, Brain, and Behavior, 18(1), e12475. 10.1111/gbb.12475

Chang, K.-J., Redmond, S. A., & Chan, J. R. (2016). Remodeling myelination: Implications for mechanisms of neural plasticity. Nature Neuroscience, 19(2), Article 2. 10.1038/nn.4200

Chase, I. D. (1982). Dynamics of Hierarchy Formation: The Sequential Development of Dominance Relationships. Behaviour, 80(3/4), 218–240.

Chase, I. D., Tovey, C., Spangler-Martin, D., & Manfredonia, M. (2002). Individual differences versus social dynamics in the formation of animal dominance hierarchies. Proceedings of the National Academy of Sciences of the United States of America, 99(8), 5744–5749. 10.1073/pnas.082104199

Chen, P., & Hong, W. (2018). Neural Circuit Mechanisms of Social Behavior. Neuron, 98(1), 16–30. 10.1016/j.neuron.2018.02.026

Chen, W.-J., Chen, H., Li, Z.-M., Huang, W.-Y., & Wu, J.-L. (2023). Acetylcholine muscarinic M1 receptors in the rodent prefrontal cortex modulate cognitive abilities to establish social hierarchy. Neuropsychopharmacology, 1–9. 10.1038/s41386-023-01785-z

Choi, G. B., Dong, H., Murphy, A. J., Valenzuela, D. M., Yancopoulos, G. D., Swanson, L. W., & Anderson, D. J. (2005). Lhx6 Delineates a Pathway Mediating Innate Reproductive Behaviors from the Amygdala to the Hypothalamus. Neuron, 46(4), 647–660. 10.1016/j.neuron.2005.04.011

Crimmins, B. E., Lingawi, N. W., Chieng, B. C., Leung, B. K., Maren, S., & Laurent, V. (2023). Basal forebrain cholinergic signaling in the basolateral amygdala promotes strength and durability of fear memories. Neuropsychopharmacology, 48(4), Article 4. 10.1038/s41386-022-01427-w

Curley, J. P. (2016). Temporal pairwise-correlation analysis provides empirical support for attention hierarchies in mice. Biology Letters, 12(5), 20160192. 10.1098/rsbl.2016.0192

De Angelis, F., Bernardo, A., Magnaghi, V., Minghetti, L., & Tata, A. M. (2012). Muscarinic receptor subtypes as potential targets to modulate oligodendrocyte progenitor survival, proliferation, and differentiation. Developmental Neurobiology, 72(5), 713–728. 10.1002/dneu.20976

Desjardins, J. K., Hofmann, H. A., & Fernald, R. D. (2012). Social Context Influences Aggressive and Courtship Behavior in a Cichlid Fish. PLOS ONE, 7(7), e32781. 10.1371/journal.pone.0032781

Fields, R. D. (2015). A new mechanism of nervous system plasticity: Activity-dependent myelination. Nature Reviews Neuroscience, 16(12), Article 12. 10.1038/nrn4023

Fields, R. D., Dutta, D. J., Belgrad, J., & Robnett, M. (2017). Cholinergic signaling in myelination. Glia, 65(5), 687–698. 10.1002/glia.23101

Friesen, C. N., Maclaine, K. D., & Hofmann, H. A. (2022). Social status mediates behavioral, endocrine, and neural responses to an intruder challenge in a social cichlid, Astatotilapia burtoni. Hormones and Behavior, 145, 105241. 10.1016/j.yhbeh.2022.105241

Gammell, M. P., Vries, H. de, Jennings, D. J., Carlin, C. M., & Hayden, T. J. (2003). David’s score: A more appropriate dominance ranking method than Clutton-Brock et al.’s index. Animal Behaviour, 66(3), 601–605.

Giardino, L., Ceccatelli, S., Zanni, M., Hökfelt, T., & Calzà, L. (1994). Regulation of VIP mRNA expression by thyroid hormone in different brain areas of adult rat. Molecular Brain Research, 27(1), 87–94. 10.1016/0169-328X(94)90188-0

Hecker, S., & Mesulam, M. M. (1994). Two types of cholinergic projections to the rat amygdala. Neuroscience, 60(2), 383–397. 10.1016/0306-4522(94)90252-6

Hobson, E. A., & DeDeo, S. (2015). Social Feedback and the Emergence of Rank in Animal Society. PLOS Computational Biology, 11(9), e1004411. 10.1371/journal.pcbi.1004411

Hong, W., Kim, D.-W., & Anderson, D. J. (2014). Antagonistic Control of Social Behaviors by Inhibitory and Excitatory Neurons in the Medial Amygdala. Cell, 158(6), 1348–1361. 10.1016/j.cell.2014.07.049

Jiang, S.-N., Cao, J.-W., Liu, L.-Y., Zhou, Y., Shan, G.-Y., Fu, Y.-H., Shao, Y.-C., & Yu, Y.-C. (2023). Sncg, Mybpc1, and Parm1 Classify subpopulations of VIP-expressing interneurons in layers 2/3 of the somatosensory cortex. Cerebral Cortex, 33(8), 4293–4304. 10.1093/cercor/bhac343

Jie, F., Yin, G., Yang, W., Yang, M., Gao, S., Lv, J., & Li, B. (2018). Stress in Regulation of GABA Amygdala System and Relevance to Neuropsychiatric Diseases. Frontiers in Neuroscience, 12, 562. 10.3389/fnins.2018.00562

Kellis, D. M., Kaigler, K. F., Witherspoon, E., Fadel, J. R., & Wilson, M. A. (2020). Cholinergic neurotransmission in the basolateral amygdala during cued fear extinction. Neurobiology of Stress, 13, 100279. 10.1016/j.ynstr.2020.100279

Kingsbury, M. A., & Wilson, L. C. (2016). The Role of VIP in Social Behavior: Neural Hotspots for the Modulation of Affiliation, Aggression, and Parental Care. Integrative and Comparative Biology, 56(6), 1238–1249. 10.1093/icb/icw122

Kljakic, O., Al-Onaizi, M., Janíčková, H., Chen, K. S., Guzman, M. S., Prado, M. A. M., & Prado, V. F. (2021). Cholinergic transmission from the basal forebrain modulates social memory in male mice. European Journal of Neuroscience, 54(6), 6075–6092. 10.1111/ejn.15400

Kuhn, S., Gritti, L., Crooks, D., & Dombrowski, Y. (2019). Oligodendrocytes in Development, Myelin Generation and Beyond. Cells, 8(11), 1424. 10.3390/cells8111424

Kuznetsova, A., Brockhoff, P. B., Christensen, R. H. B., & Jensen, S. P. (2020). lmerTest: Tests in Linear Mixed Effects Models (Version 3.1-3) [Computer software]. https://cran.r-project.org/web/packages/lmerTest/index.html

Kwon, J.-T., Ryu, C., Lee, H., Sheffield, A., Fan, J., Cho, D. H., Bigler, S., Sullivan, H. A., Choe, H. K., Wickersham, I. R., Heiman, M., & Choi, G. B. (2021). An amygdala circuit that suppresses social engagement. Nature, 593(7857), Article 7857. 10.1038/s41586-021-03413-6

Langmead, B., & Salzberg, S. L. (2012). Fast gapped-read alignment with Bowtie 2. Nature Methods, 9(4), 357–359. 10.1038/nmeth.1923

Lee, W., Dwortz, M. F., Milewski, T. M., Champagne, F. A., & Curley, J. P. (2022). Social status mediated variation in hypothalamic transcriptional profiles of male mice. Hormones and Behavior, 142, 105176. 10.1016/j.yhbeh.2022.105176

Lehmann, M. L., Weigel, T. K., Elkahloun, A. G., & Herkenham, M. (2017). Chronic social defeat reduces myelination in the mouse medial prefrontal cortex. Scientific Reports, 7, 46548. 10.1038/srep46548

Li, H., Jin, G., Zhu, P., Zou, L., Shi, J., Yi, X., Zhang, X., Tian, M., & Qin, J. (2014). Upregulation of Lhx8 increase VAChT expression and ACh release in neuronal cell line SHSY5Y. Neuroscience Letters, 559, 184–188. 10.1016/j.neulet.2013.11.047

Lohman, B. K., Weber, J. N., & Bolnick, D. I. (2016). Evaluation of TagSeq, a reliable low-cost alternative for RNAseq. Molecular Ecology Resources, 16(6), 1315–1321. 10.1111/1755-0998.12529

Martin, M. (2011). Cutadapt removes adapter sequences from high-throughput sequencing reads. EMBnet.Journal, 17(1), 10–12. 10.14806/ej.17.1.200

Maruska, K. P. (2014). Social regulation of reproduction in male cichlid fishes. General and Comparative Endocrinology, 207, 2–12. 10.1016/j.ygcen.2014.04.038

Maruska, K. P. (2015). Social Transitions Cause Rapid Behavioral and Neuroendocrine Changes. Integrative and Comparative Biology, 55(2), 294–306. 10.1093/icb/icv057

Maruska, K. P., & Fernald, R. D. (2010). Behavioral and physiological plasticity: Rapid changes during social ascent in an African cichlid fish. Hormones and Behavior, 58(2), 230–240. 10.1016/j.yhbeh.2010.03.011

Maruska, K. P., & Fernald, R. D. (2013). Social Regulation of Male Reproductive Plasticity in an African Cichlid Fish. Integrative and Comparative Biology, 53(6), 938–950. 10.1093/icb/ict017

Meyer, E., Aglyamova, G. V., & Matz, M. V. (2011). Profiling gene expression responses of coral larvae (Acropora millepora) to elevated temperature and settlement inducers using a novel RNA-Seq procedure. Molecular Ecology, 20(17), 3599–3616. 10.1111/j.1365-294X.2011.05205.x

Milewski, T. M., Lee, W., Champagne, F. A., & Curley, J. P. (2022). Behavioural and physiological plasticity in social hierarchies. Philosophical Transactions of the Royal Society B: Biological Sciences, 377(1845), 20200443. 10.1098/rstb.2020.0443

Milewski, T. M., Lee, W., Young, R. L., Hofmann, H. A., & Curley, J. P. (2025). Rapid changes in plasma corticosterone and medial amygdala transcriptome profiles during social status change reveal molecular pathways associated with a major life history transition in mouse dominance hierarchies. PLOS Genetics, 21(1), e1011548. 10.1371/journal.pgen.1011548

Mineur, Y. S., Fote, G. M., Blakeman, S., Cahuzac, E. L. M., Newbold, S. A., & Picciotto, M. R. (2016). Multiple Nicotinic Acetylcholine Receptor Subtypes in the Mouse Amygdala Regulate Affective Behaviors and Response to Social Stress. Neuropsychopharmacology, 41(6), Article 6. 10.1038/npp.2015.316

Mineur, Y. S., Mose, T. N., Blakeman, S., & Picciotto, M. R. (2018). Hippocampal α7 nicotinic ACh receptors contribute to modulation of depression-like behaviour in C57BL/6J mice. British Journal of Pharmacology, 175(11), 1903–1914. 10.1111/bph.13769

Mineur, Y. S., Mose, T. N., Maibom, K. L., Pittenger, S. T., Soares, A. R., Wu, H., Taylor, S. R., Huang, Y., & Picciotto, M. R. (2022). ACh signaling modulates activity of the GABAergic signaling network in the basolateral amygdala and behavior in stress-relevant paradigms. Molecular Psychiatry, 27(12), Article 12. 10.1038/s41380-022-01749-7

Mineur, Y. S., Obayemi, A., Wigestrand, M. B., Fote, G. M., Calarco, C. A., Li, A. M., & Picciotto, M. R. (2013). Cholinergic signaling in the hippocampus regulates social stress resilience and anxiety-and depression-like behavior. Proceedings of the National Academy of Sciences, 110(9), 3573– 3578. 10.1073/pnas.1219731110

Montero-Pedrazuela, A., Fernández-Lamo, I., Alieva, M., Pereda-Pérez, I., Venero, C., & Guadaño-Ferraz, A. (2011). Adult-Onset Hypothyroidism Enhances Fear Memory and Upregulates Mineralocorticoid and Glucocorticoid Receptors in the Amygdala. PLoS ONE, 6(10), e26582. 10.1371/journal.pone.0026582

Mori, T., Yuxing, Z., Takaki, H., Takeuchi, M., Iseki, K., Hagino, S., Kitanaka, J., Takemura, M., Misawa, H., Ikawa, M., Okabe, M., & Wanaka, A. (2004). The LIM homeobox gene, L3/Lhx8, is necessary for proper development of basal forebrain cholinergic neurons. European Journal of Neuroscience, 19(12), 3129–3141. 10.1111/j.0953-816X.2004.03415.x

O’Connell, L. A., & Hofmann, H. A. (2011). The Vertebrate mesolimbic reward system and social behavior network: A comparative synthesis. Journal of Comparative Neurology, 519(18), 3599–3639. 10.1002/cne.22735

Okada, K., Nishizawa, K., Kobayashi, T., Sakata, S., Hashimoto, K., & Kobayashi, K. (2021). Different cholinergic cell groups in the basal forebrain regulate social interaction and social recognition memory. Scientific Reports, 11(1), 13589. 10.1038/s41598-021-93045-7

Okuda, H., Tatsumi, K., Makinodan, M., Yamauchi, T., Kishimoto, T., & Wanaka, A. (2009). Environmental enrichment stimulates progenitor cell proliferation in the amygdala. Journal of Neuroscience Research, 87(16), 3546–3553. 10.1002/jnr.22160

Patel, A. J., Hayashi, M., & Hunt, A. (1987). Selective persistent reduction in choline acetyltransferase activity in basal forebrain of the rat after thyroid deficiency during early life. Brain Research, 422(1), 182–185. 10.1016/0006-8993(87)90556-7

Petrulis, A. (2020). Chapter 2—Structure and function of the medial amygdala. In J. H. Urban & J. A. Rosenkranz (Eds.), Handbook of Behavioral Neuroscience (Vol. 26, pp. 39–61). Elsevier. 10.1016/B978-0-12-815134-1.00002-7

Phatak, J., Lu, H., Wang, L., Zong, H., May, C., Rouault, M., Qutaish, M., Zhang, B., Ma, X.-J., & Anderson, C. (2022). THE RNAscope MULTIPLEX IN SITU HYBRIDIZATION TECHNOLOGY ENABLES THE INCORPORATION OF SPATIAL MAPPING AND CONFIRMATION OF GENE SIGNATURES INTO SINGLE CELL RNA SEQUENCING WORKFLOWS.

Poggi, G., Albiez, J., & Pryce, C. R. (2022). Effects of chronic social stress on oligodendrocyte proliferation-maturation and myelin status in prefrontal cortex and amygdala in adult mice. Neurobiology of Stress, 18, 100451. 10.1016/j.ynstr.2022.100451

Prado, V. F., Martins-Silva, C., de Castro, B. M., Lima, R. F., Barros, D. M., Amaral, E., Ramsey, A. J., Sotnikova, T. D., Ramirez, M. R., Kim, H.-G., Rossato, J. I., Koenen, J., Quan, H., Cota, V. R., Moraes, M. F. D., Gomez, M. V., Guatimosim, C., Wetsel, W. C., Kushmerick, C.,… Prado, M. A. M. (2006). Mice Deficient for the Vesicular Acetylcholine Transporter Are Myasthenic and Have Deficits in Object and Social Recognition. Neuron, 51(5), 601–612. 10.1016/j.neuron.2006.08.005

Purger, D., Gibson, E. M., & Monje, M. (2016). Myelin plasticity in the central nervous system. Neuropharmacology, 110, 563–573. 10.1016/j.neuropharm.2015.08.001

Raam, T., & Hong, W. (2021). Organization of neural circuits underlying social behavior: A consideration of the medial amygdala. Current Opinion in Neurobiology, 68, 124–136. 10.1016/j.conb.2021.02.008

Raymaekers, S. R., & Darras, V. M. (2017). Thyroid hormones and learning-associated neuroplasticity. General and Comparative Endocrinology, 247, 26–33. 10.1016/j.ygcen.2017.04.001

Ribeiro, F. M., Black, S. A. G., Prado, V. F., Rylett, R. J., Ferguson, S. S. G., & Prado, M. A. M. (2006). The “ins” and “outs” of the high-affinity choline transporter CHT1. Journal of Neurochemistry, 97(1), 1–12. 10.1111/j.1471-4159.2006.03695.x

Robinson, M. D., & Oshlack, A. (2010). A scaling normalization method for differential expression analysis of RNA-seq data. Genome Biology, 11(3), R25. 10.1186/gb-2010-11-3-r25

Saddoris, M. P., Gallagher, M., & Schoenbaum, G. (2005). Rapid Associative Encoding in Basolateral Amygdala Depends on Connections with Orbitofrontal Cortex. Neuron, 46(2), 321–331. 10.1016/j.neuron.2005.02.018

Sams, E. C. (2021). Oligodendrocytes in the aging brain. Neuronal Signaling, 5(3), NS20210008. 10.1042/NS20210008

Segerson, T. P., Lam, K. S. L., Cacicedo, L., Minamitani, N., Fink, J. S., Lechan, R. M., & Reichlin, S. (1989). THYROID HORMONE REGULATES VASOACTIVE INTESTINAL PEPTIDE (VIP) mRNA LEVELS IN THE RAT ANTERIOR PITUITARY GLAND. Endocrinology, 125(4), 2221–2223. 10.1210/endo-125-4-2221

Snyder-Mackler, N., Sanz, J., Kohn, J. N., Brinkworth, J. F., Morrow, S., Shaver, A. O., Grenier, J.-C., Pique-Regi, R., Johnson, Z. P., Wilson, M. E., Barreiro, L. B., & Tung, J. (2016). Social status alters immune regulation and response to infection in macaques. Science, 354(6315), 1041–1045. 10.1126/science.aah3580

So, N., Franks, B., Lim, S., & Curley, J. P. (2015). A Social Network Approach Reveals Associations between Mouse Social Dominance and Brain Gene Expression. PLOS ONE, 10(7), e0134509. 10.1371/journal.pone.0134509

Storey, J. D., & Tibshirani, R. (2003). Statistical significance for genomewide studies. Proceedings of the National Academy of Sciences of the United States of America, 100(16), 9440–9445. 10.1073/pnas.1530509100

Taborsky, B., & Oliveira, R. F. (2012). Social competence: An evolutionary approach. Trends in Ecology & Evolution, 27(12), 679–688. 10.1016/j.tree.2012.09.003

Tomioka, T., Shimazaki, T., Yamauchi, T., Oki, T., Ohgoh, M., & Okano, H. (2014). LIM Homeobox 8 (Lhx8) Is a Key Regulator of the Cholinergic Neuronal Function via a Tropomyosin Receptor Kinase A (TrkA)-mediated Positive Feedback Loop *. Journal of Biological Chemistry, 289(2), 1000–1010. 10.1074/jbc.M113.494385

Vasudevan, N., Morgan, M., Pfaff, D., & Ogawa, S. (2013). Distinct behavioral phenotypes in male mice lacking the thyroid hormone receptor α1 or β isoforms. Hormones and Behavior, 63(5), 742–751. 10.1016/j.yhbeh.2013.03.015

Walker, D. M., Zhou, X., Cunningham, A. M., Ramakrishnan, A., Cates, H. M., Lardner, C. K., Peña, C. J., Bagot, R. C., Issler, O., Van der Zee, Y., Lipschultz, A. P., Godino, A., Browne, C. J., Hodes, G. E., Parise, E. M., Torres-Berrio, A., Kennedy, P. J., Shen, L., Zhang, B., & Nestler, E. J. (2022). Crystallin Mu in Medial Amygdala Mediates the Effect of Social Experience on Cocaine Seeking in Males but Not in Females. Biological Psychiatry, 92(11), 895–906. 10.1016/j.biopsych.2022.06.026

Williamson, C. M., Franks, B., & Curley, J. P. (2016). Mouse Social Network Dynamics and Community Structure are Associated with Plasticity-Related Brain Gene Expression. Frontiers in Behavioral Neuroscience, 10. 10.3389/fnbeh.2016.00152

Williamson, C. M., Klein, I. S., Lee, W., & Curley, J. P. (2019). Immediate early gene activation throughout the brain is associated with dynamic changes in social context. Social Neuroscience, 14(3), 253–265. 10.1080/17470919.2018.1479303

Williamson, C. M., Romeo, R. D., & Curley, J. P. (2017). Dynamic changes in social dominance and mPOA GnRH expression in male mice following social opportunity. Hormones and Behavior, 87, 80–88. 10.1016/j.yhbeh.2016.11.001

Winslow, J. T., & Camacho, F. (1995). Cholinergic modulation of a decrement in social investigation following repeated contacts between mice. Psychopharmacology, 121(2), 164–172. 10.1007/BF02245626

Xin, W., & Chan, J. R. (2020). Myelin plasticity: Sculpting circuits in learning and memory. Nature Reviews. Neuroscience, 21(12), 682–694. 10.1038/s41583-020-00379-8

