## Supplemental Figures and Tables for "Rapid changes in cholinergic signaling, myelination and thyroid signaling pathway gene expression in amygdala subnuclei in response to social status maintenance and reorganization"

**Supplemental Figure 1:** Housing systems for social groups in **A)** social reorganization and **B)** vivarium paradigms.
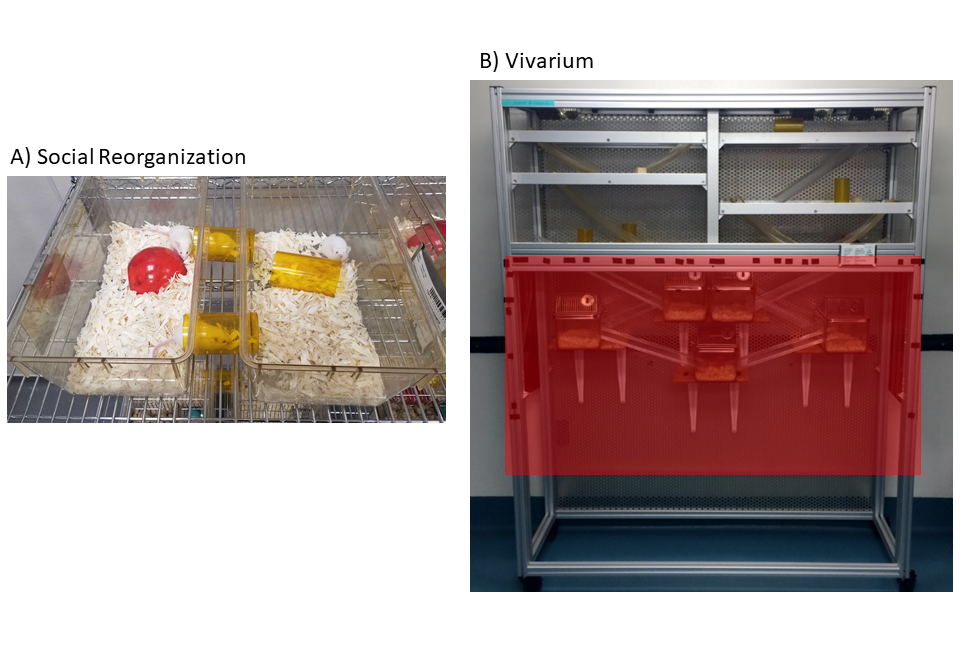

**Supplemental Figure 2: A)** Pre-Reorganized and **B)** Post- Reorganization sociomatrices of wins and losses for each cohort. Each value represents the total number of wins by the individual in each row against the individual in each column. The degree of redness represents the frequency of wins.

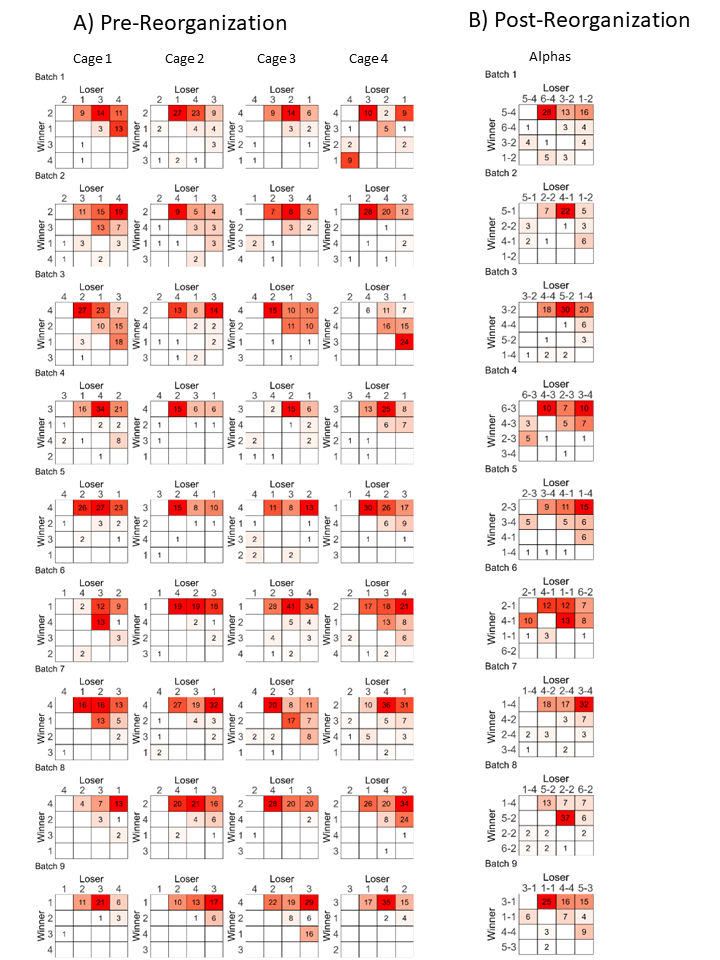

**Supplemental Figure 3: A**) Median David’s score with IQR for each social rank for all animals pre-reorganization. Gray lines represent individual animals within each cage/hierarchy. **B)** Boxplots showing median and IQR of body mass by pre-reorganization social rank at the beginning of social observation on day 1 of group housing and **C)** on day 11 of group housing. Points represent individual animals. Letters indicate which groups are significantly different from each other (p<0.05).

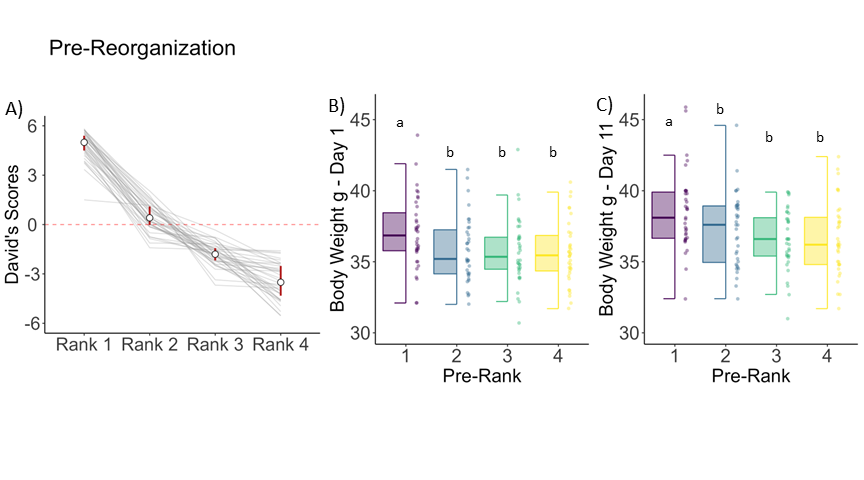

**Supplemental Figure 4: A)** Median David’s score with IQR of each social rank for males post-reorganization. Gray lines represent individual animals within each cage/hierarchy (N=9 hierarchies). Boxplots showing median and IQR of body mass on **B)** day 1 and **C)** day 11 of group housing by post-reorganization social rank. **D)** Rates of aggression given by each alpha male in their pre-reorganization cage by post-reorganization social rank **E)** There is a significant negative correlation between post-reorganization David’s Score and total aggression in each alpha male’s cage prior to reorganization.

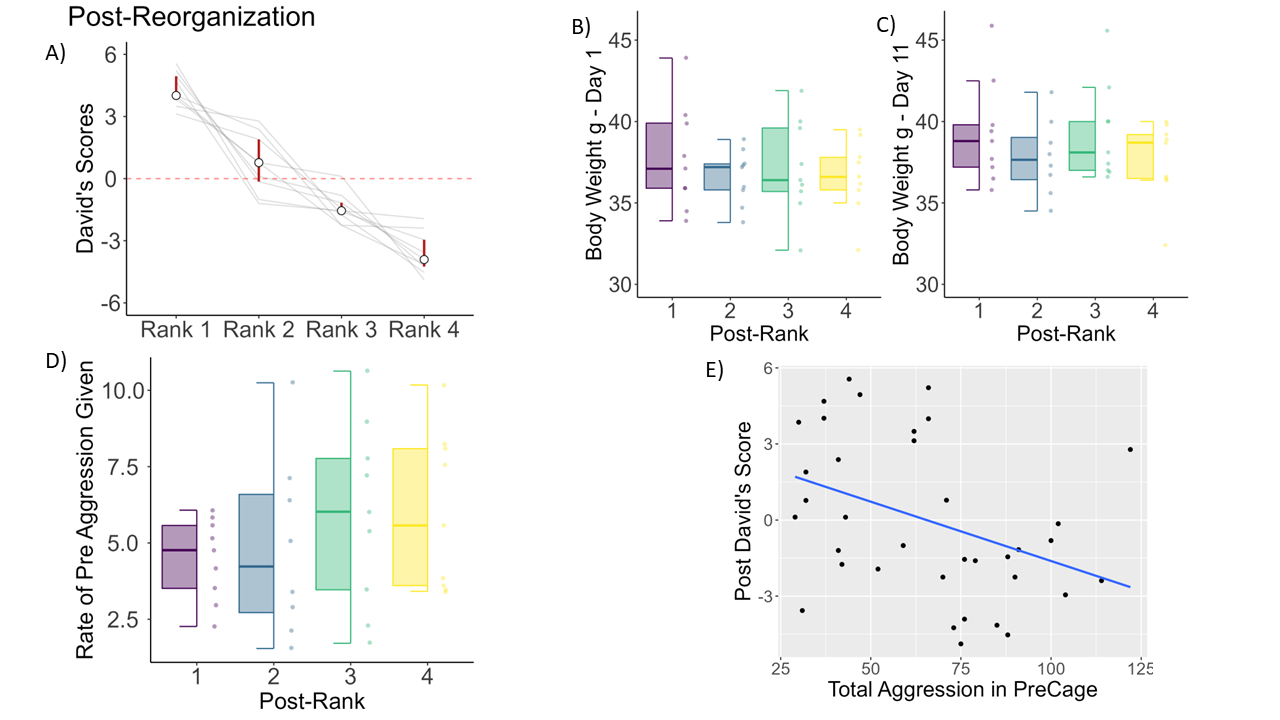

**Supplemental Figure 5** Representative images of mRNA expression of A) *crym* B) *mybpc1*, C) *mog*, D) *mbp*, E) *chat*, F) *slc5a7* in the BLA in descending (DES) versus dominant (DOM) males.

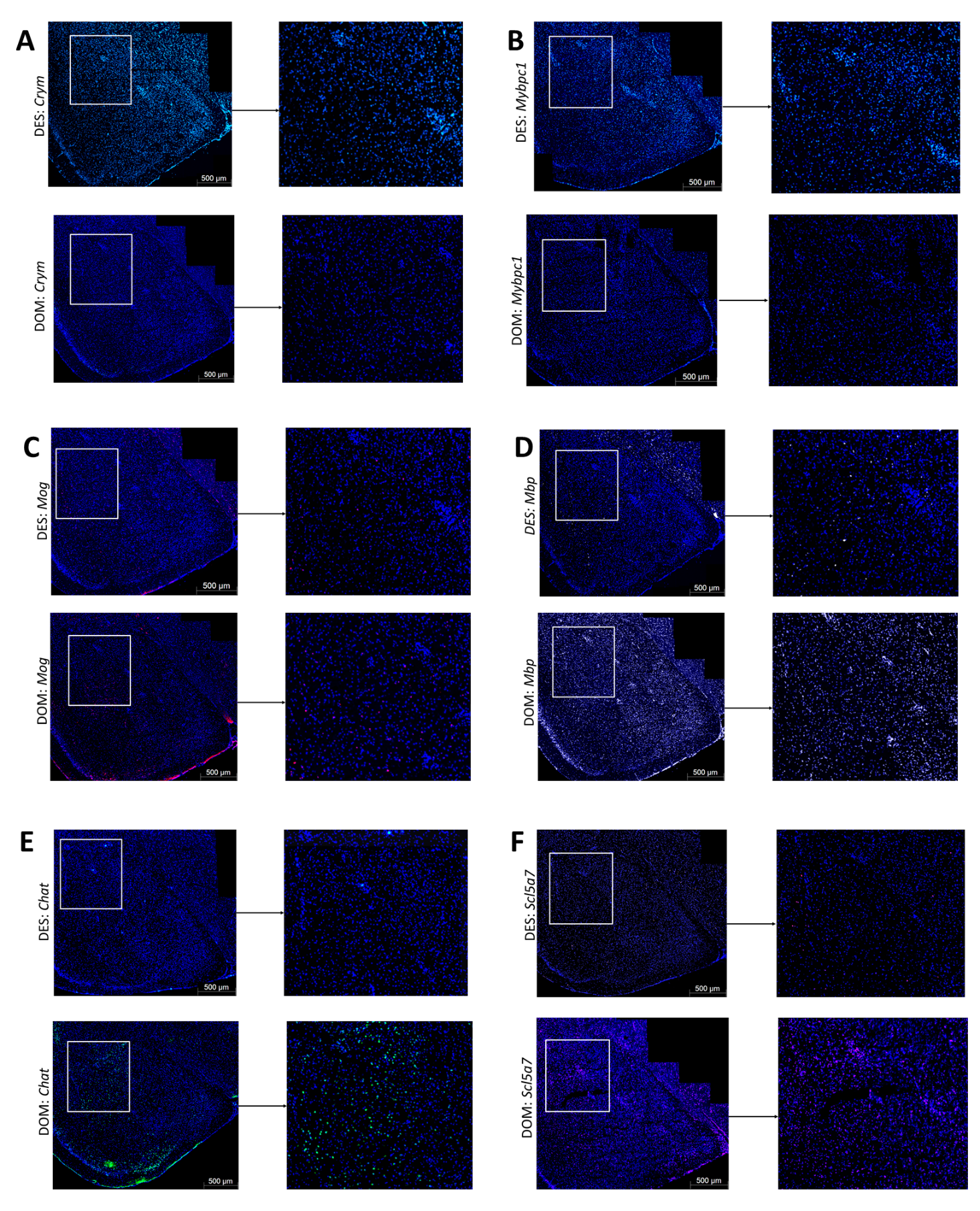

**Supplemental Table 1:** Log2 fold changes and eFDRs of cholinergic, myelination, and thyroid related genes. Log2FC = log2 fold change, eFDR = significance level. Bolded text show eFDR < 0.05.

|  | DES vs. DOM  70 min | | DES vs. DOM  25 hr | | SUB vs. DOM  stable | | SUBDOM vs. DOM stable | |
| --- | --- | --- | --- | --- | --- | --- | --- | --- |
| Gene symbol | Log2FC | eFDR | Log2FC | eFDR | Log2FC | eFDR | Log2FC | eFDR |
| Chrm1 | **0.44** | **0.013** | 0.01 | 0.909 | 0.00 | 0.966 | 0.0 | 0.892 |
| Gna11 | **0.32** | **0.005** | 0.05 | 0.146 | 0.08 | 0.626 | 0.01 | 0.740 |
| Rgs2 | 0.029 | 0.817 | -0.02 | 0.652 | -0.11 | 0.193 | **-0.20** | **0.020** |
| Ache | **-0.51** | **0.015** | -0.08 | 0.139 | **-0.14** | **0.049** | **-0.16** | **0.031** |
| Chrna2 | -0.52 | 0.212 | 0.08 | 0.502 | **-0.64** | **0.009** | -0.36 | 0.112 |
| Slc5a7 | **-1.56** | **0.004** | -0.19 | 0.284 | **-0.79** | **0.003** | **-0.66** | **0.012** |
| Chrm2 | **-1.59** | **<0.001** | **-0.43** | **0.036** | -0.21 | 0.144 | -0.13 | 0.368 |
| Chat | **-1.67** | **0.047** | -0.95 | 0.011 | -0.07 | 0.866 | -0.27 | 0.538 |
| Gbx1 | **-1.75** | **0.026** | - | - | -0.74 | 0.065 | **-1.08** | **0.014** |
| Slc10a4 | **-2.55** | **0.003** | -0.51 | 0.100 | **-0.87** | **0.011** | **-1.30** | **<0.001** |
| Slc18a3 | **-3.07** | **0.001** | - | - | - | - | - | - |
| Lhx8 | **-3.76** | **0.004** | -053 | 0.103 | **-0.91** | **0.022** | **-1.59** | **<0.001** |
| Arhgef10 | **-0.80** | **<0.001** | 0.02 | 0.859 | -0.17 | 0.183 | -0.10 | 0.401 |
| Cntn2 | **-0.80** | **<0.001** | **-0.186** | **0.004** | -0.13 | 0.128 | **-0.18** | **0.036** |
| Tspan2 | **-1.11** | **<0.001** | **-0.30** | **0.001** | -0.15 | 0.241 | **-0.29** | **0.024** |
| Opalin | **-1.14** | **0.002** | 0.05 | 0.634 | 0.03 | 0.827 | -0.15 | 0.299 |
| Myrf | **-1.14** | **0.001** | -0.09 | 0.312 | -0.27 | 0.111 | 0.01 | 0.931 |
| Cnp | **-1.17** | **<0.001** | **-0.22** | **0.020** | -0.04 | 0.734 | **-0.27** | **0.044** |
| Plp1 | **-1.31** | **0.002** | **-0.29** | **0.032** | -0.10 | 0.538 | -0.28 | 0.116 |
| Mbp | **-1.40** | **<0.001** | **-0.30** | **0.039** | -0.03 | 0.843 | -0.28 | 0.072 |
| Mag | **-1.41** | **0.001** | **-0.33** | **0.013** | -0.15 | 0.405 | -0.38 | 0.468 |
| Bcas1 | **-1.42** | **<0.001** | **-0.32** | **0.033** | -0.14 | 0.390 | -0.25 | 0.137 |
| Mal | **-1.45** | **<0.001** | **-0.28** | **0.053** | 0.01 | 0.969 | -0.33 | 0.656 |
| Mobp | **-1.59** | **<0.001** | **-0.38** | **0.036** | -0.08 | 0.686 | -0.29 | 0.176 |
| Lpar1 | **-1.65** | **<0.001** | **-0.40** | **0.038** | 0.15 | 0.380 | -0.20 | 0.280 |
| Mog | **-1.76** | **<0.001** | -0.34 | 0.074 | -0.24 | 0.222 | **-0.56** | **0.009** |
| Mybpc1 | **2.95** | **0.012** | **1.59** | **0.026** | 1.24 | 0.093 | 1.3 | 0.079 |
| Crym | **1.81** | **0.004** | 0.39 | 0.151 | **0.86** | **<0.001** | **0.87** | **0.002** |
| Vip | **1.78** | **0.008** | **0.33** | **0.007** | 0.54 | 0.167 | 0.65 | 0.107 |
| Trhr | **0.90** | **0.025** | **0.38** | **0.012** | **0.44** | **0.009** | **0.43** | **0.013** |
| Thrb | **0.58** | **0.004** | **0.15** | **0.033** | **0.24** | **0.036** | 0.21 | 0.066 |
| Thra | **0.29** | **<0.001** | 0.06 | 0.190 | **0.07** | **0.037** | 0.00 | 0.896 |
| Aldh1a3 | -0.05 | 0.78 | -0.02 | 0.771 | -0.11 | 0.644 | 0.05 | 0.703 |
| Aldh1a1 | -0.89 | 0.034 | -0.25 | 0.192 | **-0.25** | **0.042** | -0.25 | 0.362 |

**Supplemental Table 2:** RNAscope mRNA targets used in each round of staining. Catalog numbers are provided in parentheses.

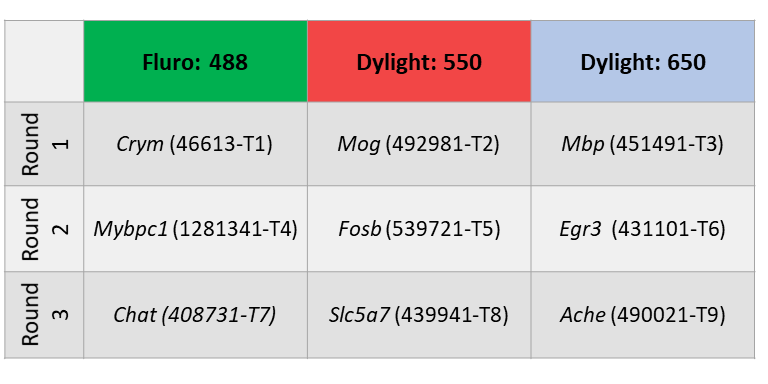
